# Molecular basis for substrate specificity of the Phactr1/PP1 phosphatase holoenzyme

**DOI:** 10.1101/2020.06.28.176040

**Authors:** Roman O. Fedoryshchak, Magdalena Přechová, Abbey Butler, Rebecca Lee, Nicola O’Reilly, Helen Flynn, Ambrosius P. Snijders, Noreen Eder, Sila Ultanir, Stéphane Mouilleron, Richard Treisman

## Abstract

PPP-family phosphatases such as PP1 have little intrinsic specificity. Cofactors can target PP1 to substrates or subcellular locations, but it remains unclear how they might confer sequence-specificity on PP1. The cytoskeletal regulator Phactr1 is a neuronally-enriched PP1 cofactor that is controlled by G-actin. Structural analysis showed that Phactr1 binding remodels PP1’s hydrophobic groove, creating a new composite surface adjacent to the catalytic site. Using phosphoproteomics, we identified numerous fibroblast and neuronal Phactr1/PP1 substrates, which include cytoskeletal components and regulators. We determined high-resolution structures of Phactr1/PP1 bound to the dephosphorylated forms of its substrates IRSp53 and spectrin αII. Inversion of the phosphate in these holoenzyme-product complexes supports the proposed PPP-family catalytic mechanism. Substrate sequences C-terminal to the dephosphorylation site make intimate contacts with the composite Phactr1/PP1 surface, which are required for efficient dephosphorylation. Sequence specificity explains why Phactr1/PP1 exhibits orders-of-magnitude enhanced reactivity towards its substrates, compared to apo-PP1 or other PP1 holoenzymes.

## INTRODUCTION

PPP family phosphatases are metalloenzymes that carry out the majority of protein serine/threonine dephosphorylation (Brautigan and Shenolikar, 2018). The three Protein Phosphatase 1 (PP1) isoforms regulate diverse cellular processes, acting in partnership with over 200 different PP1-interacting proteins (PIPs). Some PIPs are PP1 substrates, but others are PP1 cofactors, which variously determine substrate specificity, subcellular targeting and/or coupling to regulatory pathways (Bollen et al., 2010; Cohen, 2002). The PP1 catalytic site lies at the intersection of three putative substrate-binding grooves (Egloff et al., 1995; Goldberg et al., 1995), and PIPs can interact both with these grooves and with other PP1 surface features. To do this they use a variety of short sequence elements, of which the best understood is the RVxF motif (Choy et al., 2014; Egloff et al., 1997; Hendrickx et al., 2009; Hurley et al., 2007; O’Connell et al., 2012; Ragusa et al., 2010; Terrak et al., 2004).

Unlike protein Ser/Thr kinases, PP1 exhibits little sequence-specificity by itself (Brautigan and Shenolikar, 2018; Miller and Turk, 2018). Moreover, no instances of PIP-induced sequence-specificity are known, although it is well established that PIPs can enhance or inhibit PP1 activity towards particular substrates (Ichikawa et al., 1996; Johnson et al., 1996). Some PIPs contain autonomous substrate-binding domains, which facilitate substrate recruitment (Boudrez et al., 2000; Choy et al., 2015), while others constrain substrate specificity by occluding PP1 surfaces such as the RVxF-binding pocket and/or substrate-binding grooves (Hirschi et al., 2010; Ragusa et al., 2010). Interestingly, several PIPs extend the PP1 substrate-binding grooves and/or significantly alter PP1 surface electrostatics, without altering the conformation of the catalytic site (O’Connell et al., 2012; Ragusa et al., 2010; Terrak et al., 2004). How this affects substrate selection remains unclear, the sequence-specificity of these PIP/PP1 holoenzymes, and how they bind substrates, have not been characterized.

The four Phosphatase and actin regulator (Phactr) proteins (Allen et al., 2004; Sagara et al., 2003) are novel PIPs that are implicated in cytoskeletal regulation in animal models (Hamada et al., 2018; Kim et al., 2007; Zhang et al., 2012) and cell culture settings (Huet et al., 2013; Wiezlak et al., 2012). The Phactrs bind G-actin via multiple RPEL motifs present in their conserved N- and C-terminal regions (Huet et al., 2013; Mouilleron et al., 2012; Sagara et al., 2009; Wiezlak et al., 2012). Their C-terminal RPEL domain overlaps the PP1 binding sequence (Allen et al., 2004; Larson et al., 2008; Sagara et al., 2003), and G-actin competes with PP1 for Phactr binding (Huet et al., 2013; Wiezlak et al., 2012). As a result, like other RPEL proteins, G-actin/Phactr interactions respond to fluctuations in actin dynamics (Diring et al., 2019; Miralles et al., 2003; Vartiainen et al., 2007). Extracellular signals, acting through the “rho-actin” signal pathway, can control Phactr/PP1 complex formation by inducing changes in cellular G-actin concentration (Wiezlak et al., 2012).

The biochemical function of Phactr/PP1 complexes has been unclear. Phactr1 and Phactr3 inhibit dephosphorylation of phosphorylase a by PP1 *in vitro* (Allen et al., 2004; Sagara et al., 2003), but Phactr4/PP1 complex formation is associated with cofilin dephosphorylation *in vivo* (Huet et al., 2013; Zhang et al., 2012). Here we show that Phactr1 confer sequence-specificity on the Phactr1/PP1 holoenzyme. We identify substrates for Phactr1/PP1, and determine the structures of Phactr1/PP1-substrate complexes. We show that efficient catalysis requires interactions between conserved hydrophobic residues in the substrate and a novel Phactr1-PP1 composite surface, which comprises a hydrophobic pocket and associated amphipathic cavity with a surrounding basic rim. These interactions allow Phactr1/PP1 to recognise its substrates 100-fold more efficiently compared with PP1 alone or the spinophilin/PP1 complex, in which the hydrophobic groove is remodelled differently.

## RESULTS

### Crystallisation of Phactr1/PP1 complexes

Previous studies have shown that Phactr1 C-terminal sequences are necessary and sufficient for interaction with PP1 (Allen et al., 2004; Wiezlak et al., 2012): they contain an RVxF-like sequence, LIRF, and can functionally substitute for the related PP1-binding domain of yeast Bni4 (Larson et al., 2008) (Figure 1A). We synthesised peptides corresponding to Phactr1(517-580) from all four Phactr family members, and measured their affinity for recombinant PP1α(7-300) using Bio-layer interferometry (BLI). Phactr1(517-580) bound with an affinity of 10.4 nM, comparable to Phactr3, while the other Phactrs bound more weakly (Figure 1B, 1C, Figure S1A). This K_d_ is similar to PIPs such as spinophilin (8.7nM) and PNUTS (9.3nM) (Choy et al., 2014; Ragusa et al., 2010), and somewhat stronger than PPP1R15A and NIPP1 (Choy et al., 2015; O’Connell et al., 2012).

**Figure 1.**
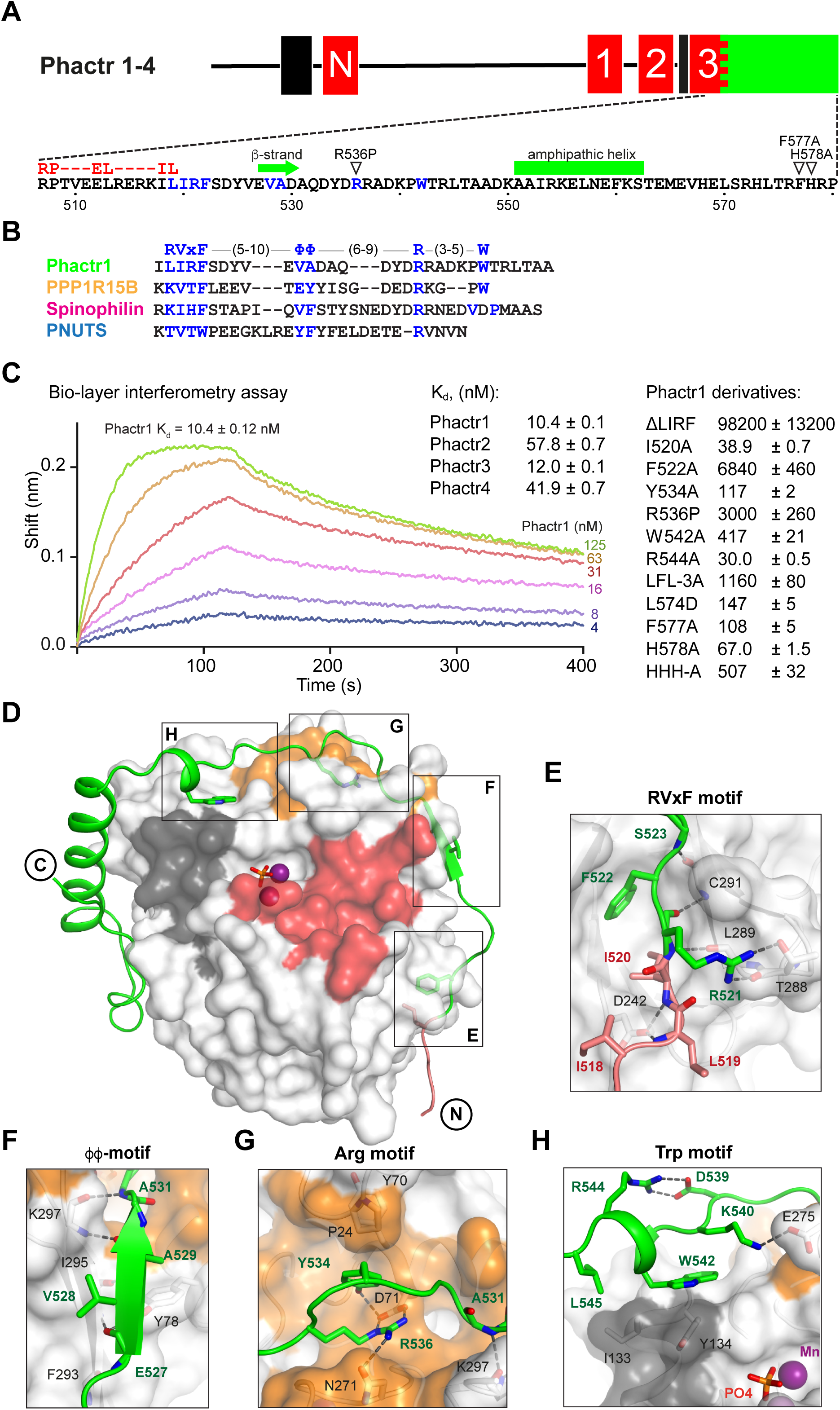
Phactr1 binds PP1 using an extended RVxF-ϕϕ-R motif. (A) Domain structure of Phactr family proteins. RPEL motifs, red; PP1-binding domain, green; nuclear localisation sequences, black. Below, Phactr1 C-terminal sequence, indicating RVxF-ϕϕ-R-W string, RPEL consensus, secondary structure elements, and mutations known to impair Phactr/PP1 interaction(Allen et al., 2004; Kim et al., 2007). (B) Structure-based alignment of Phactr1 PP1-binding sequences with other PP1 cofactors. (C) Bio-layer interferometry assay of PP1α(7-300) binding to Phactr1(517-580), its derivatives, and analogous sequences from Phactr2-4 (data are means ± SD, n = 3). (D) Structure of the Phactr1(516-580)/PP1α(7-300) complex Phactr1, green ribbon, with RPEL motif in red, RVxF-ϕϕ-R-W sidechains as sticks; PP1, white surface, with acidic groove in red, hydrophobic groove in dark grey and C-terminal groove in gold; manganese ions, purple spheres; phosphate, orange sticks. (E-H) Detail of the individual interactions, with important residues highlighted (Phactr1, green bold; PP1, black). For comparison with other PIPs, see Figure S2A.

We determined the structures of purified Phactr(507-580)/PP1α(7-300) (1.9Å) and Phactr1(516-580)/PP1α(7-300) (1.94Å), which crystallised at pH 8.5 and pH 5.25 respectively (Figure 1D, S1B). In the pH5.25 structure, the Phactr1 sequences C-terminal to residue 567 adopted an extended conformation, perhaps owing to protonation of the three C-terminal histidines, which was poorly resolved (Figure S1B). As will be described below, the pH8.5 structure is adopted by Phactr1(516-580)/PP1α(7-300) when substrates occupy the PP1 active site; it is also supported by BLI data, and is therefore likely to exist at physiological pH. The conformation of PP1 is virtually identical to that seen in other PP1 complexes, such as spinophilin/PP1 (RMS: 0.23 Å over 255 Cα atoms) (Choy et al., 2014; Ragusa et al., 2010). Its catalytic site contains two manganese ions and a presumed phosphate anion, as do other PP1 structures (Figure S1C)(Egloff et al., 1995; Goldberg et al., 1995).

### Phactr1 binds PP1 through an extended RVxF-ϕϕ-R-W string

Phactr1 residues 516-542 wrap around PP1, occluding its C-terminal groove, and covering 2260 Å^2^ of solvent-accessible surface (Figure 1D, Figure S1D), contacting PP1 in a strikingly similar way to spinophilin, PNUTS and PPP1R15B (Chen et al., 2015; Choy et al., 2014; Ragusa et al., 2010). These contacts include a non-canonical RVxF motif (Egloff et al., 1997; Hendrickx et al., 2009), a ΦΦ motif (O’Connell et al., 2012), an Arg motif(Choy et al., 2014), and a previously unrecognised Trp motif (Figure 1E-H; Figure S2A). The RVxF sequence LIRF(519-522) is critical for Phactr1/PP1 complex formation (Figure 1E, Figure S2A). Its deletion decreased binding affinity ∼10^4^-fold, while the I520A and F522A mutations reduced it 4-fold and ∼650-fold respectively (Figure 1C). The RVxF residue L519 also makes contacts with G-actin in the trivalent G-actin*Phactr1 RPEL domain complex (Mouilleron et al., 2012), explaining why PP1 and G-actin binding to Phactr proteins is mutually exclusive (Figure S2B) (Wiezlak et al., 2012).

The other contacts also contribute to PP1 binding affinity. The ϕϕ motif, EVAA(527-530), adds a β-strand to PP1 β-strand β14, extending one of PP1’s two central β-sheets (Figure 1F). The Phactr1 Arg motif contacts the PP1 C-terminal groove, with R536 forming a bidentate salt bridge with PP1 D71, and a hydrogen bond with N271. This is stabilised by an intrachain cation-π interaction with Phactr1 Y534, which also hydrogen bonds with PP1 D71, and makes hydrophobic contacts with P24 and Y70 (Figure 1G, Figure S2A). Substitution of R536 by proline, as in the mouse Phactr4 “humdy” mutation (Kim et al., 2007), reduced Phactr1 binding affinity >300-fold, while Y534A reduced it >10-fold (Figure 1C). The Trp motif, W542, which is constrained by a salt bridge between R544 and D539, makes hydrophobic contacts with PP1 I133 and Y13 (Figure 1H), similar to those in the PPP1R15B and spinophilin PP1 complexes (Chen et al., 2015; Ragusa et al., 2010) (Figure S2A). Although W542A reduced PP1 binding affinity ∼40 fold, R544A reduced it ∼3 fold (Figure 1C).

### Phactr1 binding creates a novel composite surface adjacent to PP1 catalytic site

The Phactr1 sequences C-terminal to the RVxF-ϕϕ-R-W string form a novel structure specific to the Phactr1/PP1 complex. Residues 545-565 include a five-turn amphipathic α-helix that abuts the PP1 hydrophobic groove: Phactr1 residues L545, I553, L557, F560 make hydrophobic contacts with PP1 I133, R132, W149, and K150, while E556 and E564 make salt bridges with PP1 R132 and K150 respectively, and K561 hydrogen bonds with PP1 S129 and D194 (Figure 2A). Phactr1 then turns, contacting the PP1 α7-α8 loop: Phactr1 H568 hydrogen bonds to PP1 P192 and Phactr1 S571, making hydrophobic contacts with PP1 M190, while Phactr1 L570 and L574 make hydrophobic contacts with PP1 M190 and I189/P196/L201 respectively. The Phactr1 C-terminus then folds back onto the amphipathic helix, stabilised by multiple intrachain interactions. These include hydrophobic contacts between Phactr1 T575, F577 and K561; hydrogen bonds between the R576 carbonyl and K561, between the H578 amide and N558, and between the R579 carbonyl and R554, and a salt bridge between the P580 carboxylate and the R554 guanidinium. The Phactr1 C-terminus also contacts PP1 through hydrogen bonds between R576 and PP1 D194, and H578 and PP1 S129 (Figure 2A).

**Figure 2.**
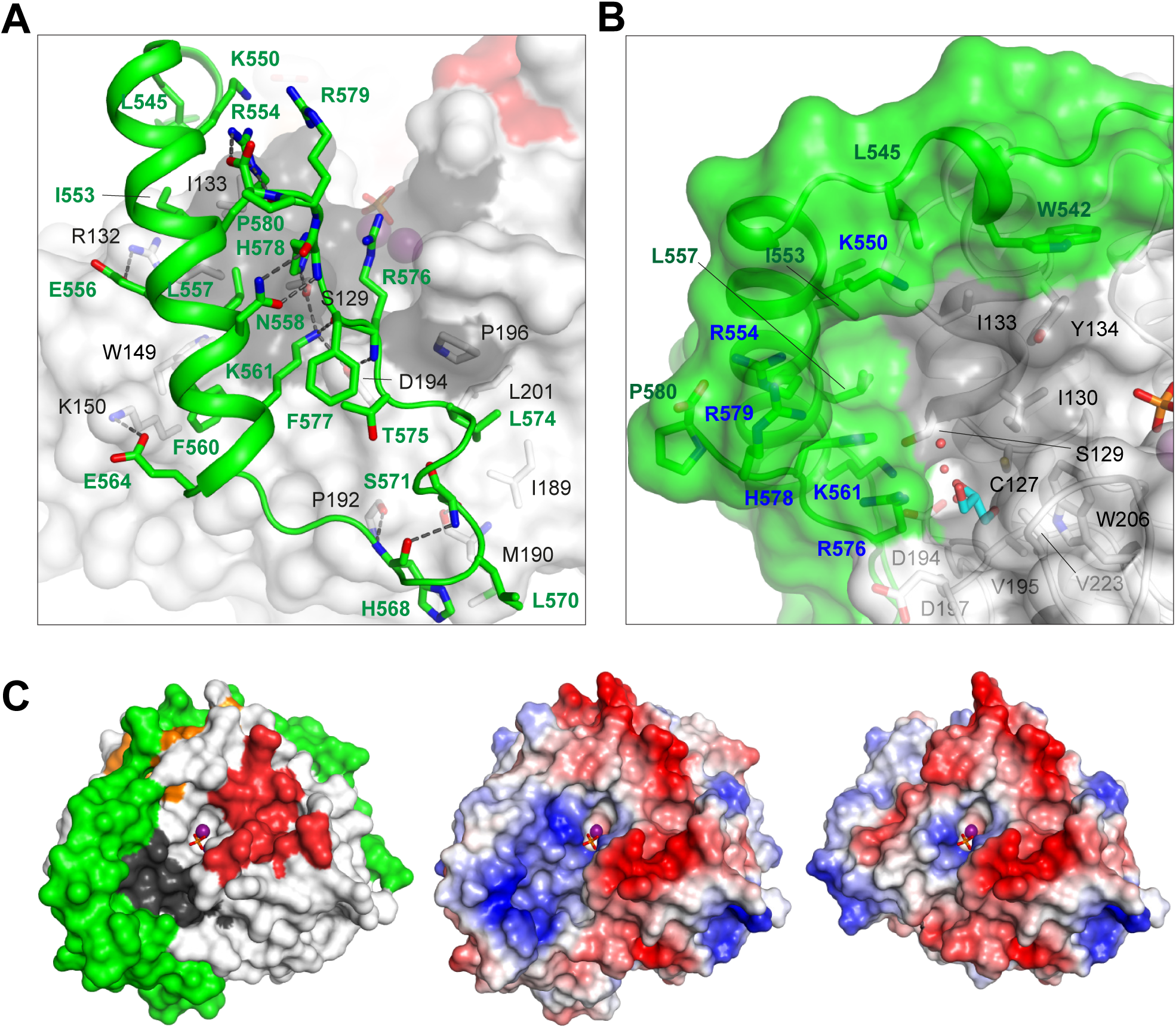
Phactr1/PP1 interaction remodels the PP1 hydrophobic groove. (A) Molecular interactions by the Phactr1 C-terminal sequences (green; residues involved in intrachain interactions or contacting PP1 are shown as sticks). (B) The novel composite surface formed by Phactr1/PP1 interaction, showing the deep hydrophobic pocket, adjacent narrow amphipathic cavity with associated waters (red spheres) and glycerol (cyan sticks), and residues constituting the Phactr1-derived basic rim (blue). (C) PP1 surface electrostatics are transformed in the Phactr1/PP1 complex. Left, surface representation of the Phactr1/PP1 complex; centre, electrostatic surface representation of Phactr1/PP1 complex. Right, electrostatic surface potential representation of PP1 (positive, blue; negative, red).

Mutagenesis experiments confirmed the importance of these interactions for Phactr1-PP1 binding (Figure 1C). A triple alanine substitution of Phactr1 L557, F560 and L574 (LFL-3A) resulted in a ∼900-fold drop in binding affinity, while the acidic substitution L574D reduced binding affinity by 17-fold. Mutations F577A and H578A reduced binding activity by 16-fold and 7-fold respectively, corroborating previous co-immunoprecipitation experiments (Allen et al., 2004; Sagara et al., 2009), and mutation of all three C-terminal histidines (HHH-A) reduced affinity >50-fold (Figure 1C).

The docking of Phactr1 C-terminal sequences across the PP1 hydrophobic groove substantially modifies its topography (Figure 2B). The positioning of the Phactr1 amphipathic helix creates a deep hydrophobic pocket comprising Phactr1 W542, L545, K550, I553, R554, L557 and H578, and PP1 I130 I133 and Y134. Adjacent to the pocket, a narrow amphipathic cavity is formed by the positioning of Phactr1 K561 and R576 across the PP1 hydrophobic groove: its polar side comprises Phactr1 K561 and R576, and PP1 D194 and S129, while its hydrophobic side is formed from PP1 residues C127, I130, V195, W206 and V223. Three water molecules and a glycerol are resolved within the cavity, whose hydrophobic side forms part of the binding site for the PP1 inhibitors tautomycetin and tautomycin (Choy et al., 2017). The new composite surface is crowned by a basic rim, formed from Phactr1 residues K550, R554, R576, H578 and R579 (Figure 2B), which radically alters the surface charge distribution (Figure 2C). Thus, Phactr1 binding profoundly transforms the surface of PP1 adjacent to its catalytic site. This transformation is distinct from that seen in the spinophilin/PP1 complex, which modifies the hydrophobic groove in a different way and leaves the PP1 surface electrostatics unchanged (Figure S2C).

### Identification of potential Phactr1 substrates

We previously showed that expression of an activated Phactr1 derivative that constitutively forms the Phactr1/PP1 complex, Phactr1^XXX^, induces F-actin rearrangements in NIH3T3 fibroblasts, provided it can bind PP1 (Wiezlak et al., 2012); indeed, expression of the Phactr1 PP1-binding domain alone is sufficient to induce such changes (Figure S3A). These observations suggest that the Phactr1/PP1 complex might dephosphorylate target proteins involved in cytoskeletal dynamics. To identify potential Phactr1/PP1 substrates, we used differential SILAC phosphoproteomics in NIH3T3 cells expressing Phactr1^XXX^. Over 3000 phosphorylation sites were quantified, and each assigned a dephosphorylation score comparing their phosphorylation with that observed in cells inducibly expressing Phactr1^XXX^ΔC, which lacks the PP1 binding sequences, or vector alone (Figure 3A; Figure S3B, Table S1A).

**Figure 3.**
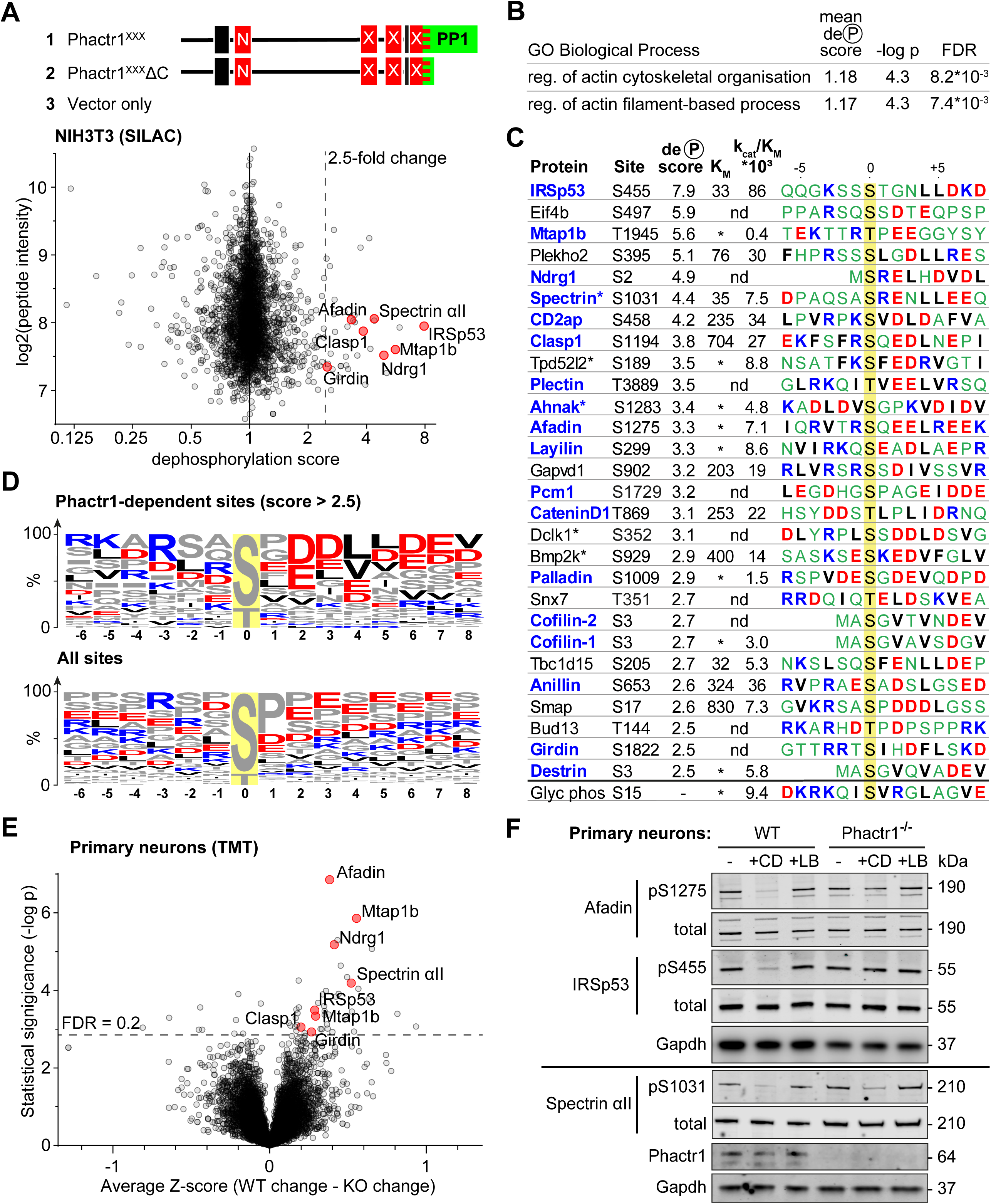
Identification of Phactr1/PP1 substrates. (A-E), NIH3T3 cells, (F,G), neuronal cells. (A) SILAC phosphoproteomics in NIH3T3 cells. NIH3T3 cell lines conditionally expressed Phactr1^XXX^ (constitutively binds PP1, not G-actin), Phactr1^XXX^ΔC (binds neither PP1 nor G-actin), or vector alone (Wiezlak et al., 2012). Forward and reverse SILAC phosphoproteomics were used to generate a dephosphorylation score, quantified below (red highlights, hits also detected as Phactr1-dependent in neurons). (B) Annotation enrichment analysis of the entire SILAC phosphoproteomics dataset for GO Biological Process terms, showing terms with dephosphorylation score > 1, FDR < 0.02. (C) Candidate Phactr1/PP1 substrates ranked by dephosphorylation score (blue, cytoskeletal structural or regulatory proteins; asterisks, proteins with multiple dephosphorylation sites). Michaelis constant (K_M_, µM), specificity constant (k_cat_/K_M_, µM*min^−1^*U^−1^) and sequence context (blue, basic; red, acidic; black, hydrophobic) are shown. (*), K_M_ could not be reliably determined; (nd), not done. (D) Amino acid frequency among phosphorylation sites with dephosphorylation score >2.5 (top) compared with all phosphorylation sites (bottom). (E) Phactr1-dependent protein dephosphorylation in cortical and hippocampal neurons treated with cytochalasin D (CD). Differential Z-score, the difference between the phosphorylation change observed in Phactr1-wildtype and Phactr1-null neurons, plotted versus statistical significance. Red highlights, peptides also observed in the NIH3T3 SILAC phosphoproteomics. (F) Validation of TMT phosphoproteomics data in primary cortical neurons treated for 30’ with CD or latrunculin B (LB). For quantitation, see Figure S3H.

Annotation enrichment analysis of the whole dataset (Cox and Mann, 2012) identified two Gene Ontology Biological Process categories that exhibited a significantly higher mean dephosphorylation score upon expression of Phactr1^XXX^: “regulation of actin filament-based process” and “regulation of actin cytoskeleton organisation” (Figure 3B, Table S1B). In keeping with this, proteins with a high dephosphorylation score included many cytoskeletal components and regulators (Figure 3C). Proline-directed sites predominated in the dataset as a whole, but those sites with dephosphorylation scores >2.5 were enriched in acidic residues at positions +2 to +7 relative to the phosphorylation site, with small hydrophobic residues enriched at positions +4 and +5, and basic residues N-terminal to it (Figure 3D). Since this sequence bias was not observed at all sites, we tested directly whether sites were Phactr1/PP1 substrates using an *in vitro* peptide dephosphorylation assay. Synthetic phosphopeptides containing the candidate sites exhibited a range of K_M_ and k_cat_/K_M_ values: for example, IRSp53 exhibited a low K_M_ and high k_cat_/K_M_; spectrin αII had a similar K_M_ but lower k_cat_/K_M_; and afadin was as reactive as spectrin αII, but with much poorer K_M_ (Figure 3C; Figure S3C). Interestingly, the four substrates with the lowest K_M_ – IRSp53, Plekho2, spectrin αII, and Tbc1d15 – all contained a leucine doublet at the +4 and +5 positions (see Discussion).

### IRSp53, afadin and spectrin αII are Phactr1/PP1 substrates

We generated phosphospecific antisera against IRSp53 pS455, spectrin αII pS1031, and afadin pS1275. Immunoblot analysis showed that in NIH3T3 cells, phosphorylation of IRSp53 S455 and afadin S1275 was substantially decreased upon expression of Phactr1^XXX^, and by serum stimulation, which activates rho-actin signalling and Phactr-family/PP1 complex formation (Miralles et al., 2003; Vartiainen et al., 2007; Wiezlak et al., 2012) (Figure S3B, S3D). Moreover, CD treatment, which disrupts G-actin interaction with RPEL proteins (Vartiainen et al., 2007; Wiezlak et al., 2012) significantly decreased phosphorylation of IRSp53 S455 and afadin S1275, while LatB treatment, which increases G-actin concentration but does not affect RPEL-actin interaction, had the opposite effect (Figure S3E). These data suggest that endogenous RPEL protein(s), presumably Phactr-family members, controls IRSp53 S455 and afadin S1275 phosphorylation in NIH3T3 cells.

To explore directly the specific involvement of Phactr1 in protein dephosphorylation we turned to neurons, which express Phactr1 at high level (Allen et al., 2004). Phactr1 mutations are associated with morphological and functional developmental defects in cortical neurons (Hamada et al., 2018), and expression of Phactr1^XXX^ induced morphological defects upon expression in cultured hippocampal neurons (Figure S3F). To assess whether Phactr1 controls phosphorylation in neurons, we analysed hippocampal and cortical neurons from wildtype and Phactr1-null animals. Phactr1-null mice are viable; they do not show any obvious developmental abnormalities, and expression of the other Phactr proteins is apparently unaffected (Figure S3G). Neurons were treated with LatB or CD to inhibit or activate RPEL proteins, and phosphorylation profiles analysed using TMT phosphoproteomics. Among ∼9000 phosphorylation sites quantified, we found 44 sites on 37 proteins that differed significantly in their response to these stimuli in Phactr1-null cells (Figure 3E, Table S2). The sequence context of these sites was similar to that of those seen in NIH3T3 cells (Figure S3H), and seven, including IRSp53 pS455, spectrin αII pS1031, and afadin pS1275, were observed in both cell types (Figure 3G, Figure S3I). In sum, these results identify multiple Phactr1 substrates in NIH3T3 cells and neurons, many of which are cytoskeletal components or regulators.

### Crystallisation of Phactr1/PP1-substrate complexes

To understand the molecular basis for substrate recognition by Phactr1/PP1, we sought to determine the structures of Phactr1/PP1-substrate complexes. We were unable to co-crystallise Phactr1/PP1 with unphosphorylated or glutamate-phosphomimetic peptides from IRSp53, afadin and spectrin αII, so we used a PP1-substrate fusion strategy similar to that used for PP5 (Oberoi et al., 2016). Fusion proteins comprising PP1(7-304) joined via a (SG)_9_ linker to unphosphorylated substrate sequences were coexpressed with Phactr1(516-580) for structural analysis. This approach allowed determination of the structures of the IRSp53(449-465) and spectrin αII (1025-1039) complexes at 1.09 Å and 1.30 Å resolution respectively (Figure 4A-C; Table 1; Figure S4A-S4D; referred to hereafter as the IRSp53 and spectrin complexes). We were unsuccessful with Tbc1d15, Plekho2, afadin, and cofilin.

**Table 1.**
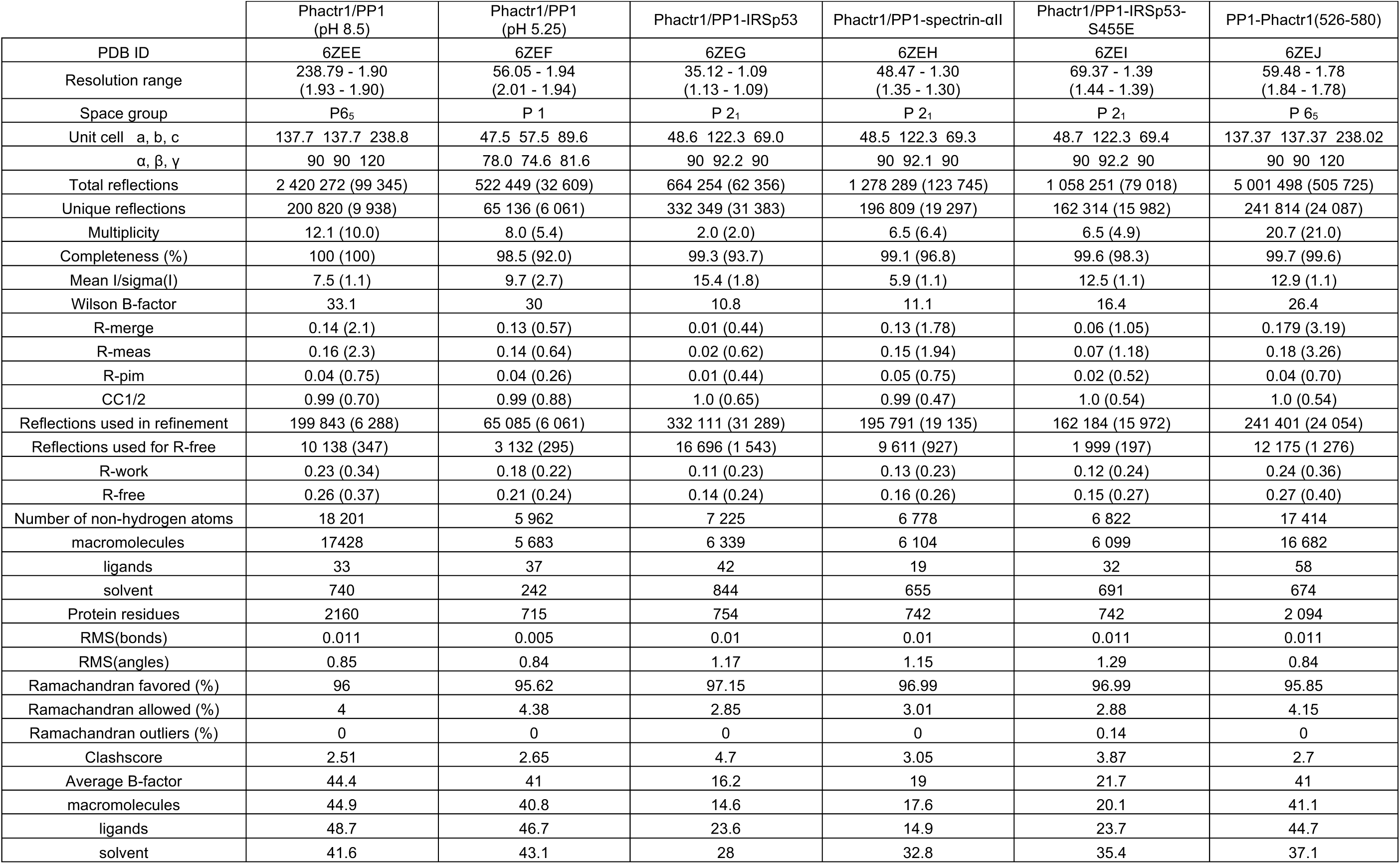
Crystallographic data and refinement statistics.

**Figure 4.**
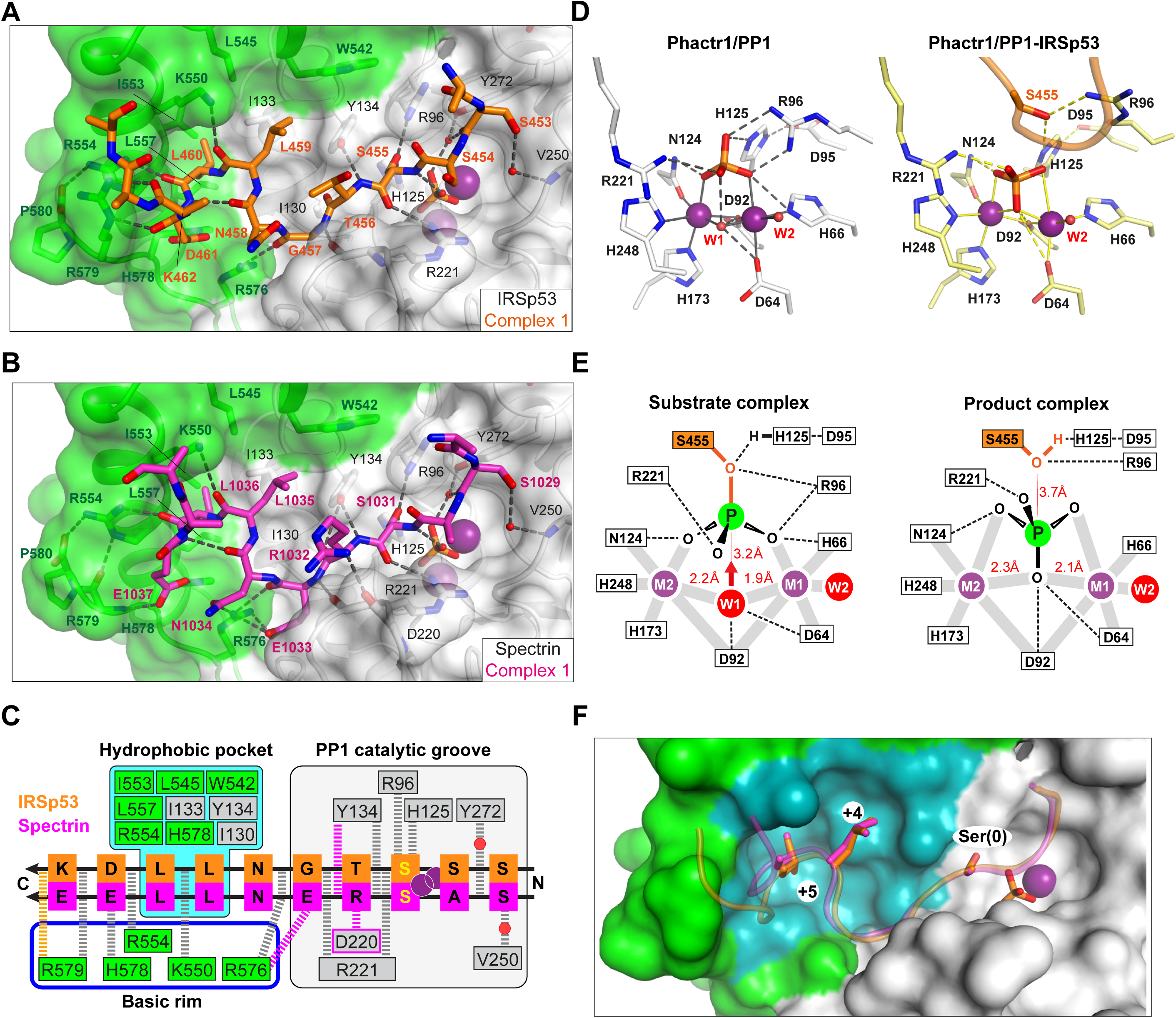
Substrate interactions with the Phactr1/PP1 holoenzyme. (A, B) Structures of (A) the Phactr1/PP1-IRSp53(449-465) and (B) the Phactr1/PP1-spectrin(1025-1039) complexes, displayed as in Figure 1, with IRSp53 and spectrin displayed in orange and magenta sticks respectively. (C) Summary of substrate interactions. Hydrogen bonds are shown as thick dashed lines: grey for both substrates; colour, for specific substrate. Composite hydrophobic surface residues are highlighted in blue (see F). (D) Inversion of the recruited phosphate. Phosphate and metal ions contacts in the Phactr1/PP1 and in Phactr1/PP1-IRSp53 structures are shown. Metal coordination bonds, solid continuous lines; hydrogen bonds, dashed lines; W1 and W2, water molecules. (E) Potential catalytic mechanism. Left, a hypothetical substrate complex, based on the Phactr1/PP1 complex, assuming that its phosphate corresponds to that of IRSp53 pS455. Right, the observed Phactr1/PP1-IRSp53 product complex. W1 and W2, water molecules; grey bars, metal coordination bonds; dashes, hydrogen bonds. Proposed nucleophilic attack by activated W1 results in phosphate inversion. (F) Docking of the SxxxLL motif (sticks) with the Phactr1/PP1 hydrophobic pocket. Phactr1/PP1 in surface representation, with the composite hydrophobic surface in light blue, and other Phactr1 and PP1 surfaces in green and white respectively.

The Phactr1/PP1-IRSp53 and Phactr1/PP1-spectrin complexes crystallised in the same spacegroup. Each asymmetric unit contains two complexes (RMSD of 0.22Å and 0.17Å over 288 Cα respectively), in which substrate-Phactr1/PP1 interactions are mostly conserved, albeit with some minor differences (Figure S4A, S4B). The substrate sequences, whose N-termini are largely unresolved, make extensive contacts across the PP1 catalytic site, then extend in a sinuous trajectory across the composite Phactr1/PP1 surface, making numerous hydrophobic and ionic contacts (Figure 4C). Like the Phactr1/PP1 holoenzyme, the complexes contain a presumed phosphate anion at the catalytic site, and thus appear to represent putative enzyme/product complexes (see below). For both substrates, the structure of Phactr1/PP1 is identical to that of the isolated Phactr1/PP1 complex (RMSD 0.31Å over 313 Cα and 0.32Å over 307 Cα, respectively).

### Phactr1/PP1-substrate interactions are opposite in polarity to those of PP5

IRSp53 and spectrin make virtually identical contacts with the PP1 catalytic cleft, predominantly via their mainchains (Figure 4C; Figure S4C). In IRSp53 complex 2, K452^IRSp53^ (−3 relative to the phosphorylation site) makes a salt bridge with PP1 D220, in the acidic groove, while water-bridged hydrogen bonds link the mainchain carbonyl and the sidechain hydroxyl of S453^IRSp53^/S1029^spectrin^(−2) to PP1 Y272 and the phosphate, and to V250 respectively. The dephosphorylated S455^IRSp53^/S1031^spectrin^(0) hydroxyl interacts with PP1 R96, H125 and the phosphate, its mainchain amide and carbonyl contacting the phosphate and PP1 R221 respectively (Figure 4A-C, Figure S4A, S4B). The phosphate is inverted compared with the holoenzyme complex, losing its contact with PP1 R96 and H125, but making contact with the S455^IRSp53^/S1031^spectrin^ hydroxyl (Figure 4D; Figure S4A-S4C). C-terminal to the dephosphorylated serine, T456^IRSp53^/R1032^spectrin^(+1) hydrogen bonds with PP1 Y134 via its mainchain amide, with R1032^spectrin^ making two additional hydrogen bonds, with PP1 Y134 via its carbonyl, and the D220 carbonyl via its sidechain.

The majority of the PP1 catalytic cleft residues that contact IRSp53 and spectrin are conserved among PPP family members, including PP5, which has been crystallized in complex with a phosphomimetic derivative of its substrate, Cdc37(S13E) (Oberoi et al., 2016). Strikingly, in that complex, the PP5 residues corresponding to PP1 R96, H125, Y134, R221, and Y272 make contacts with Cdc37(S13E) analogous to those seen in the Phactr1/PP1-substrate complexes, despite the fact that Cdc37(S13E) docks in the PP5 catalytic cleft in the opposite orientation to IRSp53 and spectrin (Figure S4E; see Discussion).

### Catalytic mechanism

It is generally accepted that the phosphate seen in many PPP family protein structures binds the active site in a similar way to the substrate phosphate (Egloff et al., 1995; Griffith et al., 1995; Mueller et al., 1993; Swingle et al., 2004), and we therefore assume that the phosphate in our Phactr1/PP1 holoenzyme structure is positioned similarly to the phosphorylated S455^IRSp53^/S1031^spectrin^ of the bound substrate (Figure 4D). Consistent with this idea, in the PP5/Cdc37(S13E) complex, the phosphomimetic glutamate sidechain carbonyl is virtually superposable on the phosphate in the Phactr1/PP1 complex (Figure S4E, S4F). The Phactr1/PP1 complex also contains a bound water, W1, presumably activated by the metal ions and PP1 D64 and D92 (Figure 4D). W1 is oriented appropriately for in-line nucleophilic attack on the substrate phosphate: this would be facilitated by protonation of the phosphoserine oxygen by the PP1 catalytic dyad H125-D95, and would result in inversion of the phosphate (Figure 4E). The structures of our Phactr1/PP1-IRSp53/spectrin complexes are consistent with their representing the resulting enzyme-product complexes, stablished through the tethering of the substrate sequences to the holoenzyme. The structures thus provide direct evidence for the in-line nucleophilic hydrolysis mechanism for PPP family phosphatases, as outlined by the Barford and Ciszak groups (Egloff et al., 1995; Swingle et al., 2004).

### Substrates make extensive contacts with the novel Phactr1/PP1 surface

C-terminal to their interaction with the catalytic cleft, both IRSp53 and spectrin follow similar trajectories from residues +2 to +7(Figure 4A-C, Figure S4C). The most striking feature of the complexes is the multiplicity of substrate contacts made between the substrate and the novel composite Phactr1/PP1 surface created by extension of the PP1 hydrophobic groove, most of which are conserved between the two substrates (Figure 4C, 4F). In both structures, the residues +3 to +6 form a β-turn, allowing the hydrophobic doublet L459-L460^IRSp53^/L1035-L1036^spectrin^ (+4/+5) to make intimate contact with the Phactr1/PP1 hydrophobic pocket, entirely burying the +5 leucine (Figure 4C, 4F). These interactions are stabilised by hydrogen bonds between the mainchain carbonyls of substrate residues +2, +4, +5 and Phactr1 R576 K550, and R554, and between the sidechain of the substrate acidic +6 residue (D461^IRSp53^/E1037^spectrin^) and Phactr1 H578 (Figure 4A-C); a hydrogen bond between the D461^IRSp53^ mainchain carbonyl and Phactr1 R579 is substituted by a hydrogen bond between the E1037^spectrin^ carboxylate and Phactr1 R579 mainchain amide. At position +2 (G457^IRSp53^/E1033^spectrin^), the mainchain amide and carbonyl make hydrogen bonds with the mainchain carbonyls of PP1 R221 and Phactr1 R576 respectively; in addition, the E1033^spectrin^ side chain spans the top of the amphipathic cavity to make an additional salt bridge with the Phactr1 R576 side chain (Figure 4B, 4C).

### Phactr1/PP1 hydrophobic pocket interactions promote catalytic efficiency

To investigate the functional significance of the Phactr1/PP1-substrate interactions seen in the structures, we tested mutated IRSp53 peptide substrates in the *in vitro* dephosphorylation assay (Figure 5A). Alanine substitutions at positions S453 (−2), S454 (−1), and T456 (+1) decreased catalytic efficiency somewhat, apparently reflecting an increased K_M_. In contrast, alanine substitution of residues L459, L460 and D461 (+4 to +6) individually increased K_M_, but had variable effects on catalytic efficiency, with L459A and D461A increasing it, and L460A reducing it (see Discussion). Pairwise combination of substitutions at positions +4 through +6 suppressed reactivity even more, with the LL459/460AA mutant being essentially unreactive. Alanine substitution at K462 (+7) and D463 (+8) either decreased or increased K_M_ with corresponding effects on catalytic efficiency, consistent with a preponderance of acidic residues at positions +6 to +8 (see Figure 3D). Upon expression in NIH3T3 cells, IRSp53(L460A) exhibited enhanced basal phosphorylation, and was less susceptible to CD-induced dephosphorylation than wildtype IRSp53, as assessed using the phospho-S455 antibody (Figure 5B). These data show that substrate interactions with the Phactr1/PP1 hydrophobic pocket are important for efficient dephosphorylation both *in vitro* and *in vivo*.

**Figure 5.**
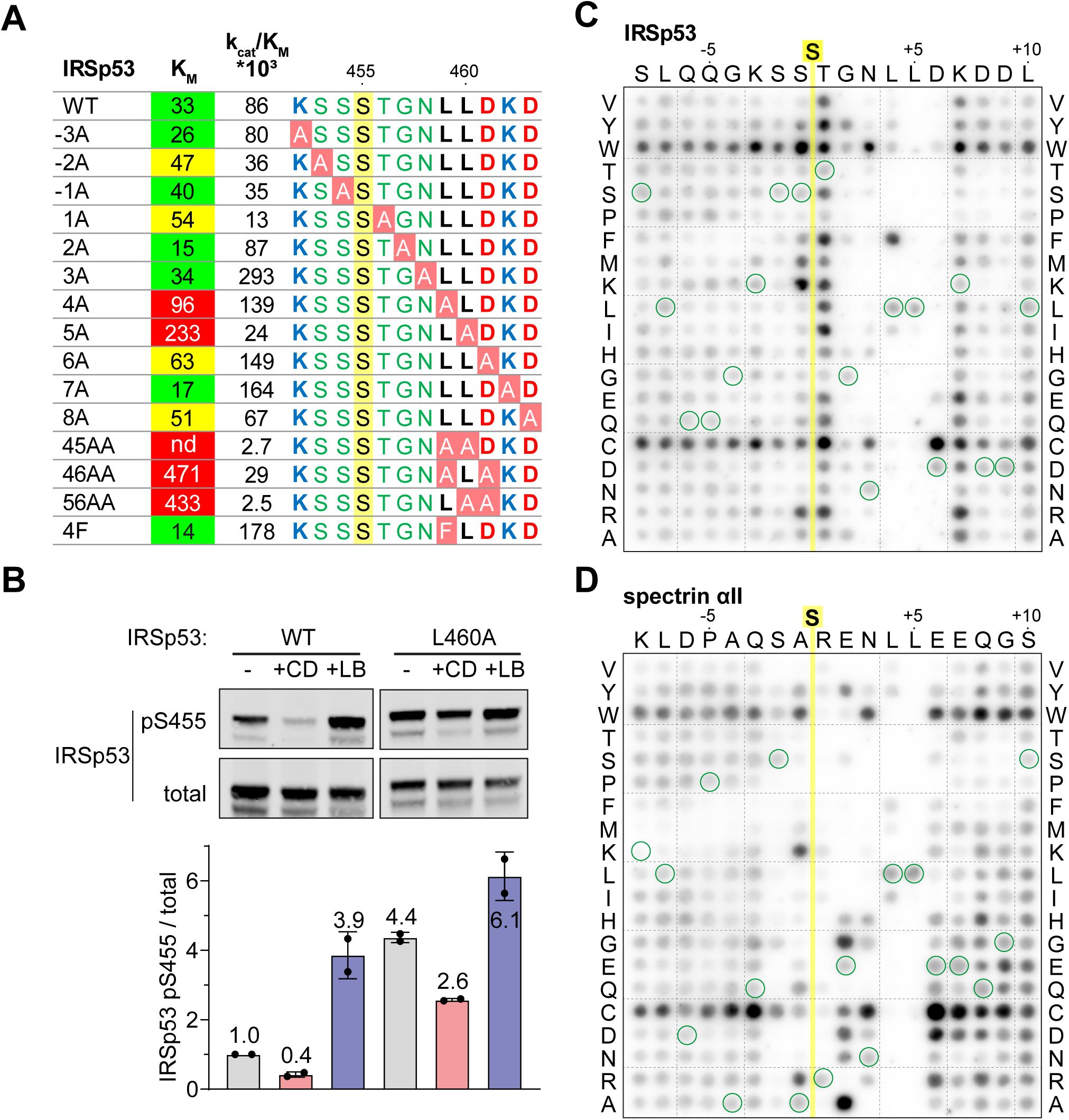
Efficient dephosphorylation involves substrate interaction with the Phactr1/PP1 composite surface. (A) Phactr1/PP1 dephosphorylation of alanine substitution derivatives of IRSp53 S455 substrate 19mer phosphopeptides. K_M_ values are highlighted: green, < 40 µM; yellow, 40-80 µM; red, >80 µM. (B) Immunoblot analysis of total IRSp53 and IRSp53 phospho-S455 levels after expression wildtype IRSp53 or IRSp53 L460A in NIH3T3 cells with 30’ CD or LB treatment as indicated. (C,D) Overlay binding affinity assay of IRSp53 (C) and spectrin αII (D). Arrays contained the variants of the wildtype sequence, in which each amino acid is systematically changes to other amino acid as indicated vertically, with wildtype sequence circled in green. Yellow line, position of the invariant unphosphorylated target serine.

The effects of substrate alanine mutations on K_M_ suggests that they affect the affinity of Phactr1/PP1-substrate interactions. To investigate substrate binding directly, we used a peptide overlay assay in which unphosphorylated substrate peptides were immobilised on membranes, and tested for their ability to recruit recombinant GST-Phactr1(516-580)/PP1(7-300) complex from solution. Since these peptides lack phosphoserine, it is likely that their binding will predominantly be determined by interactions outside the catalytic site. Only peptides exhibiting a low K_M_ IRSp53, spectrin αII, Tbc1d15 and Plekho2 exhibited detectable binding under the conditions of the assay (Figure S5; see Figure 3C). To probe the contribution of individual residues to binding we used IRSp53 and spectrin arrays in which each substrate residue was systematically changed to every other amino acid (Figure 5C, 5D). Tryptophan and cysteine substitutions at non-critical residues led to a general increase in binding affinity, and we therefore did not attempt to interpret these substitutions. Analysis of both arrays implicated residues −2 to +7 in substrate binding affinity. Positions +4 and +5 displayed the strongest selectivity, for hydrophobic residues and leucine respectively: indeed, phenylalanine substitution of IRSp53 L459 (+4), increased binding affinity and catalytic efficiency (Figure 5A). In contrast, the sequence dependence at other positions was strongly context-dependent. In the vicinity of the target residue, IRSp53 binding was relatively unaffected by N-terminal variation, while T456 (+1) was suboptimal, with basic or hydrophobic residues being preferred; in contrast, spectrin binding was strongly selective at N-terminal positions, with a preference for basic residues at −1 and +1 and serine at −2. Similarly, IRSp53 exhibited a strong preference for acidic residues at position +6, perhaps reflecting the presence of the suboptimal neighbouring basic residue, K462 (+7), which was not the case for spectrin. Taken together with the kinetics data, these results show that interactions with the Phactr1/PP1 hydrophobic pocket are a critical determinant of Phactr1/PP1 substrate recognition.

### Variable spacing between the dephosphorylation site and hydrophobic residues

In IRSp53, the dephosphorylated residue S455 is flanked by three other potential phosphoacceptors, S453, S454, and T456. We considered the possibility that strong interactions with the Phactr1/PP1 hydrophobic pocket might also allow efficient dephosphorylation of these residues. That this might be the case was suggested by our structural analysis of a phosphomimetic fusion construct, in which the S455 is substituted by glutamate, at 1.39Å resolution (Phactr1/PP1-IRSp53(S455E)). Strikingly, in this complex, the phospomimetic glutamate (S455E) does not replace phosphate at the catalytic site. Instead, IRSp53 T456 occupies the position corresponding to S455 in the wildtype complex, with a recruited phosphate positioned as in the wildtype complex, and the glutamate (S455E) in effect occupies position −1, pointing away from the catalytic site (Figure 6A). Nevertheless, the critical contacts between IRSp53(S455E) L459/L460 and the Phactr1/PP1 hydrophobic pocket are maintained, extending the intervening sequence G457-N458-L459 (Figure 6B). Consistent with this, both wildtype IRSp53 pT456 and IRSp53 E455/pT456 were effective substrates for Phactr1/PP1, exhibiting 3-5 fold higher K_M_, but only two-fold lower catalytic efficiency, while IRSp53 pS453 and pS454 were very poor substrates, exhibiting 10-30 fold lower catalytic efficiency (Figure 6C). Thus, Phactr1/PP1 substrates can tolerate either 3 or 4 residues between the target residue and the hydrophobic residue that engages the Phactr1/PP1 hydrophobic pocket (see Discussion).

**Figure 6.**
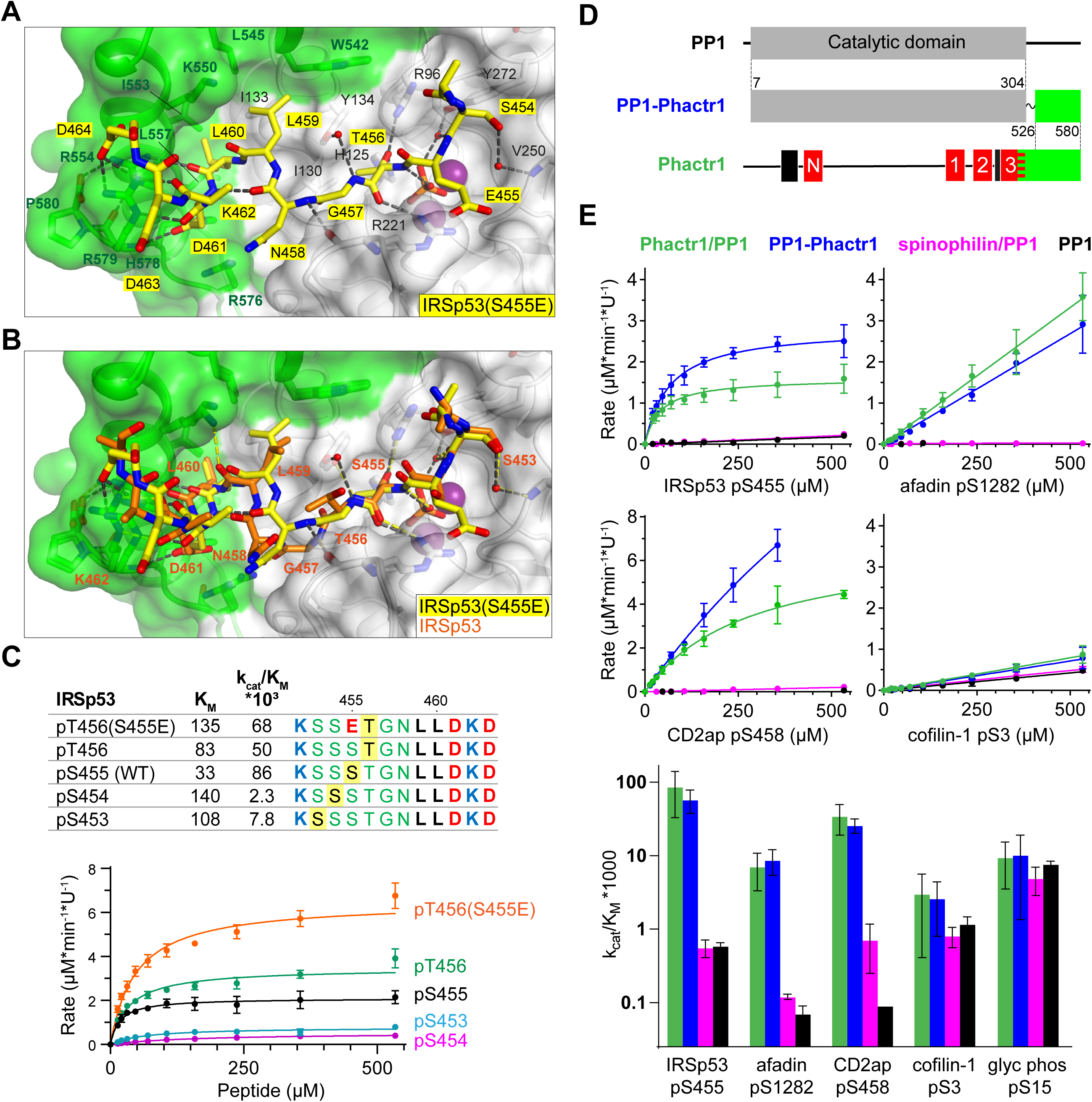
Flexible substrate interactions and substrate specificity of the Phactr1/PP1 complex. (A, B) Flexibility in target serine-hydrophobic pocket binding residue spacing, illustrated by (A) structure of the Phactr1/PP1-IRSp53(S455E) complex, compared with (B) the Phactr1/PP1-IRSp53 wildtype complex. (C) Phactr1/PP1 dephosphorylation of derivatives of IRSp53 peptides carrying phosphate at different locations, highlighted in yellow. Phosphatase activity data is shown below (data are mean ± SD, WT n= 15, others n= 2-6) (D) Schematic of the PP1-Phactr1 fusion protein PP1-Phactr1(526-580). (E) Top, phosphatase activity data for the indicated substrates and enzymes. Bottom, relative catalytic efficiencies for the different substrates (data are mean ± SD, n= from 1 to 15).

### Interaction with Phactr1 confers substrate specificity on PP1

Since the composite hydrophobic surface of the Phactr1/PP1 holoenzyme plays an important role in substrate binding and catalytic efficiency, we next investigated to what extent interaction with Phactr1 confers substrate specificity on PP1. We were particularly interested to test whether the high K_M_ and lower binding affinity of Phactr1/PP1 substrates such as afadin and cofilin might be associated with an increased promiscuity in their reactivity with different PP1 complexes. In addition to comparing Phactr1/PP1 with apo-PP1, we therefore also compared it with the spinophilin/PP1α complex (Figure S2A, S6A; hereafter spinophilin/PP1) (Ragusa et al., 2010). As discussed above, spinophilin also interacts with PP1 through an extended RVxF-ϕϕ-R-W motif, but remodels the hydrophobic groove differently, and does not change PP1 surface electrostatics (Ragusa et al., 2010) (Figure S2C). We also assessed the substrate specificity of a fusion protein in which the Phactr1 residues 526-580 were fused via a short SG linker to PP1 residue 304, solved at 1.78 Å resolution (Figure 6D). This fusion protein lacks the RVxF-PP1 interaction critical for formation of the authentic Phactr1/PP1 complex, but nevertheless generates a composite hydrophobic surface virtually identical to that seen in the Phactr1/PP1 holoenzyme (RMS 0.25 Å over 2395 atoms; Figure S6B).

We tested a series of Phactr1/PP1 substrate phosphopeptides with decreasing catalytic efficiency and increasing K_M_, and analogous phosphopeptides from glycogen phosphorylase and GluR1, substrates of the GM/PP1 and spinophilin/PP1 complexes (Hu et al., 2007; Ragusa et al., 2010). Compared with apo-PP1 and spinohilin/PP1, Phactr1/PP1 exhibited 100- to 400-fold greater catalytic efficiency against its substrates IRSp53 pS455, CD2ap pS458, and afadin pS1275 (Figure 6E). However, while Phactr1/PP1 dephosphorylated cofilin pS3 with a low catalytic efficiency comparable to afadin pS1275, this was only 2-fold enhanced compared with PP1 or spinophilin/PP1 (Figure 6E; see Discussion). Thus at least some Phactr1 substrates with high K_M_, are likely to be substates for multiple different PP1 complexes (see Discussion). Phactr1 did not enhance the activity of PP1 against the GM/PP1 substrate glycogen phosphorylase pS15, or the spinophilin/PP1 target GluR1 pS863 (Figure 6E, Figure S6C). The PP1-Phactr1 fusion protein behaved similarly to the Phactr1/PP1 complex, indicating that RVxF interactions are not essential for specificity (Figure 6E). We found that spinophilin did not enhance PP1 activity against the short GluR1 pS863 peptide (Figure S6C), although it is active against a longer GluR1 fragment (Ragusa et al., 2010), suggesting that substrate specificity is not directed by its modified hydrophobic groove. Taken together with the results in the preceding sections, these results show that the composite surface formed by interaction with Phactr1 is responsible for the substrate specificity of the Phactr1/PP1 holoenzyme.

## DISCUSSION

Here we elucidate the mechanism by which interaction of the RPEL protein Phactr1 with PP1 defines the substrate specificity of the Phactr1/PP1 holoenzyme. The Phactr1 C-terminal residues remodel the PP1 hydrophobic groove to forming a new composite hydrophobic pocket and an adjacent amphipathic cavity, topped by a basic rim. Having identified Phactr1/PP1 substrates, we showed that they make extensive contacts with the remodelled PP1 hydrophobic groove, which are required for efficient catalysis, and allow definition of a core dephosphorylation motif. The sequence conservation between Phactr family members suggest that other Phactr-family/PP1 holoenzymes will have similar specificity. The substrates exhibit >100-fold enhanced reactivity with Phactr1/PP1, when compared with apo-PP1, or the spinophilin/PP1 complex, whose remodelled hydrophobic groove is different. Our results shed new light onto the role of PIPs in substrate recognition by PPP-family phosphatase complexes.

Phactr1 embraces PP1 using an extended RVxF-ϕϕ-R-W sequence string used by several other PIPs. We exploited the close approach of the Phactr1 sequences to the PP1 C-terminus to generate a PP1-Phactr1 fusion, which has a similar specificity to intact Phactr1/PP1 holoenzyme. This approach is likely to be applicable to PNUTS, PPP1R15A/B and spinophilin/neurabin, since these proteins all use an RVxF-ϕϕ-R-W string to bind PP1 in a strikingly similar way to Phactr1. The overlap of the RVxF motif with RPEL3 explains why G-actin competes with PP1 to bind Phactr1 and Phactr4, allowing the formation of the Phactr1/PP1 complex to be controlled by fluctuations in actin dynamics (Huet et al., 2013; Wiezlak et al., 2012). Although both G-actin and PP1 bind Phactr1 with nanomolar affinity, their binding sites are overlapping rather than superposable, which should facilitate their exchange.

### Phactr1/PP1 substrates include many cytoskeletal proteins and regulators

We used proteomic approaches coupled with Phactr1 overexpression and inactivation studies to show that Phactr1/PP1 dephosphorylates multiple target proteins involved in cytoskeletal structures or regulation. This supports previous studies showing that cytoskeletal phenotypes result from Phactr1 and Phactr4 mutations (Hamada et al., 2018; Kim et al., 2007; Zhang et al., 2012), and are induced by overexpression of Phactr1 and Phactr4 derivatives that constitutively associate with PP1 (Huet et al., 2013; Wiezlak et al., 2012). Inspection of the top-scoring Phactr1/PP1 substrates shows that acidic residues are over-represented C-terminal to the target phosphorylation site, with hydrophobic residues are preferred at positions +4/+5.

Signal-regulated dephosphorylation of actin regulators by Phactr/PP1 complexes provides a new perspective on cytoskeletal regulation. For example IRSp53 was previously characterised as a Cdc42/Rac effector that controls F-actin assembly at membrane protrusions (Scita et al., 2008). The Phactr1/PP1 target residue pS455 characterised here is one of four negatively-acting putative AMPK sites that recruit 14-3-3 proteins (Cohen et al., 2011; Kast and Dominguez, 2019; Robens et al., 2010) and our data indicate another of these sites, S367, is also a Phactr1/PP1 target (Table S1). Rho-actin signalling thus provides an additional positively acting signal input to IRSp53, probably controlling interaction of the neighbouring SH3 domain with its effectors. Interestingly, the Phactr1/PP1 sites in spectrin αII and girdin are also adjacent to SH3 and SH2 domains respectively (Lin et al., 2014)(Figure 3C; Table S1). Our data also confirm that the actin depolymerising factors cofilin and destrin are also dephosphorylated (and activated) by Phactr1, in agreement with previous studies of Phactr4 (Huet et al., 2013). Since Phactr proteins are inactivated by G-actin, their activation of cofilin potentially provides a feedback loop that couples F-actin severing to decreased G-actin level, but more work is required to establish this.

Phactr1 is enriched at the post-synaptic density (PSD) (Allen et al., 2004), and is required for neuronal migration and arborization (Hamada et al., 2018). We identified multiple Phactr1-dependent substrates in hippocampal neurons. Several of these, including IRSp53 and spectrin αII, are dephosphorylated during long-term potentiation (LTP) (Li et al., 2016), and IRSp53-null mice exhibit deficits in hippocampal learning and memory (Bobsin and Kreienkamp, 2016; Kang et al., 2016). Moreover, dephosphorylation of cofilin is implicated in early-stage dendritic spine remodelling, along with G-actin itself (Bosch et al., 2014; Lei et al., 2017). These data suggest that rho-actin signalling to Phactr1 and the resulting protein dephosphorylations may contribute to synaptic plasticity, and indeed in humans Phactr1 mutations cause the infantile seizure condition West syndrome (Hamada et al., 2018). There may be multiple targets for rho-actin signalling to RPEL proteins in this setting, as the RPEL protein ArhGAP12 (Diring et al., 2019) also influences dendritic spine morphology (Ba et al., 2016). Future work will focus on the functional significance of neuronal rho-actin signalling to Phactr/PP1 substrates.

### Determinants of Phactr1/PP1 substrate recognition

We used a PP1-substrate peptide fusion strategy to characterise Phactr1/PP1-substrate interactions. In our structures, the substrate sequences are not phosphorylated, but a phosphate is recruited to the catalytic cleft: they thus appear to represent enzyme/product complexes, presumably stabilized by virtue of the protein fusion. The inversion of the phosphate relative to its orientation in the Phactr1/PP1 holoenzyme provides persuasive support for the in-line nucleophilic attack model for PPP phosphatases proposed by others (Egloff et al., 1995; Swingle et al., 2004). Substrate interaction with the catalytic cleft, which does not change its conformation, is predominantly mediated by mainchain interactions with residues conserved among the PPP family. We were surprised to see that Phactr1/PP1 substrates dock with the catalytic cleft in the opposite orientation to that previously seen in a complex between PP5 and a phosphomimetic substrate derivative (Oberoi et al., 2016). However, the clear sequence bias observed in Phactr1/PP1 substrate sequences suggests that the orientation seen in our IRSp53 and spectrin αII complexes is strongly preferred.

The most striking aspect of substrate recognition is the role played by the composite surface generated by the close apposition of the Phactr1 extreme C-terminal sequences and the PP1 hydrophobic groove. This creates a new hydrophobic pocket, into which the substrate +4/+5 hydrophobic residues are inserted, and an adjacent amphipathic cavity, which appears less important, making no specific contacts with IRSp53, and being only partially occupied by spectrin αII. The preference for acidic residues C-terminal to the dephosphorylation site presumably reflects the basic electrostatics associated with the novel composite Phactr1/PP1 surface. Biochemical studies suggest that the additional binding energy provided by from substrate interactions with the hydrophobic pocket is responsible for the enhanced catalytic rate of Phactr1/PP1 compared with apo-PP1 or other PP1 complexes. According to this view, it might be expected that substrates with higher K_M_ might be more promiscuous in their interaction with PP1 complexes. Cofilin pS3 might be such a substrate: it exhibits similar catalytic efficiency to afadin pS1275, but has little preference for Phactr1/PP1 over apo-PP1 or spinophilin/PP1. However, high binding affinities might limit catalytic efficiency by slowing product dissociation, a situation perhaps exemplified by spectrin. PP1 inhibitors generally lack specificity because they target the catalytic site (Zhang et al., 2013), and our findings suggest propose that targeting the composite Phactr1/PP1 surface may be a good strategy to create specific Phactr/PP1 inhibitors.

A striking consequence of the winding trajectory of the substrate following the catalytic site, and its strong interaction with the hydrophobic pocket, is that variation in the number of residues between the two can be tolerated. For example, IRSp53 pT456, with only four residues to the pocket-bound L460, has comparable kinetic properties to the *bona fide* IRSp53 pS455. We speculate that Phactr1/PP1 substrates such as afadin pS1275, where the sole hydrophobic residue is at +4 relative to the target residue, may well represent such “stretched” substrates. This flexibility complicates the unambiguous definition of the dephosphorylation consensus site, and further studies will be required to produce a structure-based alignment of the substrates that we have identified.

We propose that Phactr/PP1 substrates can be defined by a hydrophobic doublet at position +4/5 or +3/4 relative to the dephosphorylation site, with leucine preferred at the distal position, embedded within acidic sequences. The simple Phactr1/PP1 core recognition sequence, S/T-x_(2-3)_-ϕ-L, is reminiscent of those seen for protein kinases (Miller and Turk, 2018), and is perhaps the first identified for PP1. Mutation of positions +4 and/or +5 in candidate Phactr/PP1 substrates will be a useful way to generate authentic constitutively phosphorylated mutants for functional studies, as an alternative to “phosphomimetic” glutamate substitution of the phosphorylated residue.

### Comparison with other substrate-specificity PIPs

The lack of structural information for PP1-substrate complexes has hitherto precluded demonstration of how PIP-dependent remodelling of the substrate-binding grooves of PP1 can influence its ability to recognise substrate. In spinophilin, which also binds PP1 via a RVxF-ϕϕ-R-W string, sequences C-terminal to is W-motif form a helix that remodels the PP1 hydrophobic groove differently (Ragusa et al., 2010). Spinophilin therefore does not detectably enhance PP1 activity towards Phactr/PP1 substrates, although the relative contributions of the spinophilin-remodelled PP1 surface, and the adjacent PDZ domain, to substrate specificity remain to be determined (Ragusa et al., 2010). In contrast, MYPT interacts with PP1 in a different way to Phactr1 and spinophilin, extending both the C-terminal and acidic grooves (Terrak et al., 2004), and it will be interesting to see how this facilitates substrate recognition. In all these complexes, PIP-PP1 interaction also acts indirectly to constrain substrate binding specificity, by occluding potential substrate-binding sites on the PP1 surface, particularly such as the RVxF binding pocket (Ragusa et al., 2010).

Our results underscore the importance of the composite Phactr1/PP1 surface in substrate recognition and specificity. Nevertheless, as with protein kinases, these interactions alone are unlikely to define substrate selection completely (Miller and Turk, 2018). Phactr-family members do not appear to contain any conserved domains that might represent autonomous substrate-binding domains, as seen in NIPP1 (Boudrez et al., 2000) and PPP1R15A (Chen et al., 2015; Choy et al., 2015; Crespillo-Casado et al., 2018). On the other hand, many PIPs act to target PP1 complexes to specific subcellular compartments, proteins or macromolecules (Cohen, 2002). Phactr-family proteins exhibit differential subcellular localisations, including the nucleus and cell membrane (Huet et al., 2013; Wiezlak et al., 2012), and Phactr1 has been shown to interact directly with the KCNT1 potassium channel (Ali et al., 2020; Fleming et al., 2016). At least for Phactr3 and Phactr4, membrane targeting involves conserved N-terminal sequences (Huet et al., 2013; Itoh et al., 2014) which overlap a G-actin controlled nuclear import signal in Phactr1. The genetic dominance of West syndrome Phactr1 mutations and Phactr4 R536P/humdy mutation also is consistent with Phactr-family proteins interacting with other cellular components in addition to PP1 (Hamada et al., 2018; Kim et al., 2007; Zhang et al., 2012). These considerations, and the coupling of Phactr1/PP1 holoenzyme formation to rho-actin signalling, make it likely that the range of substrates controlled by Phactr1/PP1 in a particular setting will reflect the state of cellular actin dynamics and subcellular localisation of the complexes as well as the direct recognition of substrate primary sequence by the holoenzyme described here.

## AUTHOR CONTRIBUTIONS

**RF** identified neuronal Phactr1 substrates, analysed proteomic data, conducted the enzymology and substrate binding studies, generated phospho-specific antisera, and devised the PP1-Phactr1 fusion protein and dephosphorylation resistant IRSp53 mutants. **MP** characterised the Phactr1/PP1 complex, identified and characterised fibroblast Phactr1/PP1 substrates, generated phospho-specific antisera, generated the Phactr1-null mouse. **MP** and **RF** conducted neuron transfections. **AB** and **RL** expressed and purified proteins and conducted crystallisations. **NO’R** synthesised peptides and arrays. **HF** and **APS** conducted proteomic analysis. **NE** and **SU** cultured primary neurons and conducted proteomic analysis. **SM** conducted biophysical studies, devised the substrate fusion strategy, and determined the structures. **RT** conceived the project, designed experiments, and wrote the paper with RF, MP, and SM.

### ACKNOWLEDGEMENTS

We thank the Crick Science Technology platforms for support and advice during this work,, Clare Watkins and Julie Bee (Biological Resources), Namita Patel, Damini Patel and Alireza Alidoust (Fermentation Facility), Graham Clark (Genomics Equipment Park), Kurt Anderson (Light Microscopy) and Phil Walker and Andrew Purkiss (Structural biology). X-ray data were collected at the Diamond Light Source on beamlines I04-1 (mx9826-17), I02 beamline (mx9826-26), I03 (mx18566-37), I24 (mx18566-38), I04 (mx18566-29) and I04-1 (mx18566-55). We thank Wolfgang Peti (Brown University) and Eunjoon Kim (KAIST) for the spinophilin and IRSp53 expression plasmids respectively, and Michael Way, Neil McDonald, Peter Parker, Nic Tapon and members of the RT group for helpful discussions throughout the course of this project and/or comments on the manuscript. This work was supported by ERC Advanced Grant 268690 to RT; by Cancer Research UK core funding until March 31^st^ 2015; and since 2015 by the Francis Crick Institute, which receives its core funding from Cancer Research UK (FC001-190, FC001-115), the UK Medical Research Council (FC001-190, FC001-115) and the Wellcome Trust (FC001-190, FC001-115).

## METHODS

### Plasmids

pET28 PP1(7-330) and pcDNA3.1 IRSp53 were gifts from Wolfgang Peti and Eunjoon Kim (KAIST, S.Korea) respectively. Other plasmids were: modified pTRIPZ (Diring et al., 2019); pEF Phactr1 and derivatives (Wiezlak et al., 2012); pGEX 6p2 (GE Healthcare); and pGRO7 (Takara). For protein expression, Phactr1(507-580) and Phactr1(516-580) sequences were expressed using pGEX-6P2. PP1-substrate chimeras were derivatives of pET28 PP1(7-330) in which PP1 sequences 7-304 were joined by a (Ser-Gly)_9_ linker to either IRSp53(449-465) (QQGKSSSTGNLLDKDDL) IRSp53(449-465)-S455E (QQGKSSETGNLLDKDDL), or spectrin αII (1025-1039) (DPAQSASRENLLEEQ). pET28 PP1-Phactr1(526-580), derived from pET28 PP1(7-330), encodes PP1(7-304)-SGSGS-Phactr1(526-580). Plasmid construction and mutagenesis used standard methods, the NEB NEBuilder HiFi DNA Assembly Cloning Kit, or the NEB Q5® Site-Directed Mutagenesis Kit. Primers are listed in Table S2.

### Protein expression and purification

Protein expression in BL21 (DE3) *E. coli* cells (Invitrogen) was with pGRO7 coexpression as described (Choy et al., 2014). Overnight pre-cultures (400ml) were grown in LB medium supplemented 1 mM MnCl_2_ and used to inoculate a 100L fermenter. After growth to OD_600_ of ∼0.5, 2 g/L of arabinose was added to induce GroEL/GroES expression. At OD_600_ ∼1, the temperature was lowered to 10°C and protein expression induced with 0.1 mM IPTG for ∼18 hours. Cells were harvested, re-suspended in fresh LB medium/1mM MnCl_2_/200 μg/ml chloramphenicol and agitated for 2h at 10°C. Harvested cells were resuspended in lysis buffer (50 mM Tris-HCl, pH 8.5, 5 mM imidazole, 700 mM NaCl, 1mM MnCl_2_, 0.1% v/v TX-100, 0.5 mM TCEP, 0.5 mM AEBSF, 15 μg/ml benzamidine and complete EDTA-free protease inhibitor tablets), lysed by French press, clarified, and stored at −80°C. Phactr1/PP1 complexes were batch-adsorbed onto glutathione-sepharose affinity matrix, and recovered by cleavage with 3C protease at 4°C overnight in 50 mM Tris-HCl, pH 8.5, 500 mM NaCl, 0.5 mM TCEP. Eluted complex was further purified via adsorption on Ni-NTA IMAC, and elution with 50 mM Tris pH 8.0, 200 mM Imidazole, 700 mM NaCl and 1mM NiCl_2_ at 4°C. Finally proteins were purified using size exclusion chromatography on a Superdex 200 26/60 column equilibrated in complex buffer (20 mM Tris-HCl pH 8.5, 0.25 M NaCl and 0.4 mM TCEP).

### Bio-layer interferometry

Bio-layer interferometry (BLI) was as described (Bertran et al., 2019), using the Octet Red 96 (ForteBio). 50 μg/ml HIS-tagged PP1α was immobilised on Nickel-coated biosensor (Ni-NTA, ForteBio), and the loaded biosensors then incubated with 0.1-10 μM Phactr1 peptides in Octet buffer (50 mM Tris pH 7.5, 500 mM NaCl, 0.5 mM TCEP, 0.1% Tween 20, 500 mg BSA/100 ml). Curve fitting, steady state analysis, and calculation of kinetic parameters were done using Octet software version 7.1 (ForteBio). For peptides used, see Table S2.

### Crystallisation and structure determination

Phactr1/PP1 in complex buffer was concentrated to 10 mg/ml and crystallised at 20°C using sitting-drop vapour diffusion. Sitting drops of 1 μl consisted of a 1:1 (vol:vol) mixture of protein and well solution. Well solutions were as follows. Phactr(507-580)/PP1α(7-300): 7.5% PEG 3350, 0.2 M MgSO_4_; Phactr1(516-580)/PP1α(7-300): 1M LiCl, 0.1 M tri-sodium citrate pH 5.25, 10 % PEG 6000; Phactr1/PP1-IRSp53(S455E): 20% PEG 3350, 0.2 M NaBr; PP1-Phactr1(526-580), 20% PEG 3350, 0.2M Potassium Citrate. Crystals appeared within 24-48 hours and reached their maximum size after 4 to 7 days, apart from Phactr(507-580)/PP1α(7-300), for which the best crystals appeared after 3-7 weeks, reaching their maximum size after 8 weeks. For Phactr1/PP1-IRSp53 and Phactr1/PP1-spectrin α II complexes, crystallisation was achieved by microseed matrix screening (D’Arcy et al., 2014) using Phactr1/PP1-IRSp53(S455E) crystals. Phactr1/PP1-IRSp53 crystallised in 20% PEG 3350, 0.2 M KSCN,0.1 M BIS-Tris propane pH 8.5; Phactr1/PP1-spectrin αII crystallised in 20% PEG 3350, 0.2 M NaI, 0.1 M BIS-Tris propane pH 8.5. Crystals appeared within a day and reached maximum size within 4 to 5 days.

All crystals were cryoprotected in well solution supplemented with 15% Glycerol + 15% ethylene glycol and flash-frozen in liquid nitrogen. X-ray data were collected at 100 K at beamlines I04-1 (mx9826-17), I02 (mx9826-26), I03 (mx18566-37), I24 (mx18566-38), I04 (mx18566-29) and I04-1 (mx18566-55) of the Diamond Light Source Synchrotron (Oxford, UK). Data collection and refinement statistics are summarized in Table 1. Data sets were indexed, scaled and merged with xia2 (Winter et al., 2013). Molecular replacement used the atomic coordinates of human PP1 from PDB 4M0V (Choy et al., 2014) in PHASER (McCoy et al., 2007). Refinement used Phenix (Adams et al., 2010). Model building used COOT (Emsley et al., 2010) with validation by PROCHECK (Vaguine et al., 1999). Figures were prepared using PYMOL (Schrodinger, 2020).

Atomic coordinates and crystallographic structure factors have been deposited in the Protein Data Bank under accession codes PDB 6ZEE, Phactr(507-580)/PP1α(7-300); PDB 6ZEF, Phactr1(516-580)/PP1α(7-300); PDB 6ZEG, Phactr1/PP1-IRSp53; PDB 6ZEH, Phactr1/PP1-spectrin; PDB 6ZEI, Phactr1/PP1-IRSp53(S455E); PDB 6ZEJ, PP1(7-304)-SGSGS-Phactr1(526-580).

### Phosphatase assays

Full assay data can be found in Table S3.

Phosphatase assays (50 µl) were performed in 96-well plates. Peptides (40 µl) were serially diluted 1.5-fold from 1mM in complex buffer (20 mM Tris-HCl pH 8.5, 250 mM NaCl, 0.4 mM TCEP). Phactr1/PP1 (0.2–3U, 10 µl) was added and after 15 min at room temperature, reactions were quenched by addition of 100 µl of Biomol Green reagent (Enzo Life Sciences) for 30 min, Absorbances at 620 nm were measured using the SpectraMax Plus 384 microplate reader and converted into phosphate concentrations of using standard curves. Rate constants were estimated in GraphPad Prizm 8 by fitting product concentration readouts to modified Michaelis-Menten equation:

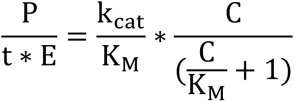

(P, product released at time t; E, Phactr1/PP1 concentration; k_cat_/K_M_, catalytic efficiency; C, initial substrate concentration; K_M_, Michaelis constant).

The activity of the Phactr1/PP1 preparation was established on the day of each experiment, using 125 µM IRSp53(447-465) pS455 peptide as standard, with Phactr1/PP1 at various concentrations. One unit of phosphatase activity was defined as the concentration of Phactr1/PP1 complex that generates 15 µM of phosphate in 15 min. To normalise the activity of different phosphatases, a para-nitrophenylphosphate (pNPP)-based assay was used. 10 µl phosphatase (200 nM) was added to 40 µl pNPP (2-fold serial dilution from 100 mM), incubated at room temperature for 15 min, then quenched with 25 µl 3 M NaOH. Para-nitrophenol product was measured by absorbance at 405 nm.

### Peptides and peptide array binding assays

Peptides were synthesized by the Francis Crick Institute Peptide Chemistry Science Technology Platform using standard techniques. Peptide arrays (5-10 nmol/spot) were synthesised on a derivatised cellulose membrane Amino-PEG_500_-UC540 using Intavis ResPep SLi automated synthesiser (Intavis Bioanalytical Instruments).

Dry membranes were blocked for 1h in 5% milk in Tris-buffered saline supplemented with 0.1% Tween-20 (TBST) with agitation, rinsed with TBST, and incubated overnight at 4°C with 10 µg/ml GST-Phactr1(516-580)/PP1(7-300) complex in TBST. After 3 10 min washes with TBST, membranes were incubated with 1:5000 HRP-conjugated anti-GST in 5% milk/TBST at room temperature for 1 h, washed 3 times with TBST, and binding revealed using SuperSignal West Pico Plus reagent (ThermoFisher) was used as chemiluminescent substrate. Images were taken using Amersham Imager 600 (GE).

### Cells, transfection and RNA analysis

Mouse NIH3T3 fibroblasts (mycoplasma-free) were maintained in DMEM (Gibco) with 10% fetal calf serum (FCS) and penicillin-streptomycin at 37°C and 5% CO_2_. Cells were transfected using Lipofectamine 2000 (Invitrogen) (150,000 cells per well, 6-well dish. For SILAC proteomics, NIH3T3 cell line pools were generated stably carrying doxycyline-inducible pTRIPZ-Phactr1 derivatives, using puromycin selection. Phactr1 expression was induced by doxycycline addition as indicated in the figure legends. Cells were maintained overnight in DMEM/0.3% FCS, and then stimulated with 15% FCS for 1 h or Cytochalasin D (CD) or latrunculin B (LatB) for 30 min. Primary rat hippocampal neurons (DIV 14) were cultured as described (Baltussen et al., 2018) and transfected using Lipofectamine 2000.

Total RNA was purified using the GenElute mammalian total RNA kit (Sigma) and cDNA synthesised using the Transcriptor First Strand cDNA Synthesis kit (Roche) with random hexamer primers. Real-time-qPCR was performed using the 7500 Fast Real-Time PCR System (Thermo Fisher Scientific) with the SYBR green reaction mix (Life Technologies). Primers are listed in Table S2. The relative abundance of target cDNA was normalised against rps16 cDNA abundance in each sample.

### Immunofluorescence and immunoblotting

Immunofluorescence microscopy in fibroblasts was performed as described (Vartiainen et al., 2007; Wiezlak et al., 2012). F-actin was detected with FITC-phalloidin (Invitrogen) and nuclei were visualised using DAPI. For immunofluorescence microscopy primary rat neurons were transfected with expression plasmids and fixed 24h later in 4% paraformaldehyde / 4% sucrose in PBS before staining with Flag and GFP antibodies. Images were taken with Leica SP5 confocal microscope with 63x (NA 1.4) oil objective. GFP was used to monitor the shape of dendritic spines, and Flag for transfected cells. An image stack with 0.5 μm z-intervals was obtained to capture a part dendritic arbour at high magnification. In each experiment, at least 10 cells per mutant were imaged and the morphology of dendritic spines was quantified blindly. Statistical analysis was performed in GraphPad Prizm 8 using Welch’s t-test function.

SDS-PAGE analysis of cell lysates and immunoblotting was performed using standard techniques; the signal was visualised and quantified using Odyssey CLx instrument (LI-COR) and the Image Studio (LI-COR) Odyssey Analysis Software.

Primary antibodies used were anti-Flag (Sigma, F7425), anti-IRSp53 (Santa Cruz, sc-50011 and Abcam, ab15697), anti-afadin (Santa Cruz, sc-74433), anti-GST (VWR, RPN1236). Secondary antibodies labelled with IRDye 800CW and IRDye 680LT were from LICOR. Rabbit anti-IRSp53 pS455, anti-afadin pS1275 and anti-spectrin pS1031 antibodies were custom-made (Covalab); for antigen sequences used see Table S2.

### SILAC phosphoproteomics in NIH3T3 cells

Cells were maintained for at least 6 passages in “heavy” (R10K8) or “light” (R0K0) DMEM medium supplemented with 10% SILAC dialysed fetal calf serum (3 kDa cutoff). Phactr1 expression was induced by doxycycline addition to 1μg/ml for 5h. Cells were harvested processed for mass spectrometry essentially as described (Pattison et al., 2016; Touati et al., 2018) with minor modifications. 1 mg each of “light” and “heavy” labelled lysates were mixed and dried. Protease digestion and phosphopeptide enrichment was done as described with minor modifications (Pattison et al., 2016; Touati et al., 2018). Peptides were dissolved in 35 μl of 1% TFA, with sonication, and fractionated on a 50 cm, 75 μm I.D. Pepmap column with elution directly into the LTQ-Orbitrap Velos. Xcalibur software was used to setup data-dependent acquisition in top10 mode. Raw mass spectrometric data were processed in MaxQuant (version 1.3.0.5) for peptide and protein identification; database search was with the Andromeda search engine against the *Mus musculus* canonical sequences from UniProtKB. Fixed modifications were set as Carbamidomethyl (C) and variable modifications set as Oxidation (M), Acetyl (Protein N-term) and Phospho (STY). The estimated false discovery rate was set to 1% at the peptide, protein, and site levels. A maximum of two missed cleavages were allowed. Phosphorylation site tables were imported into Perseus (v1.4.0.2) for analysis. Contaminants and reverse peptides were cleaned up from the Phosphosites (STY).

Dephosphorylation score was defined as log_2_(dephosphorylation score) = 0.25*[log_2_(2^H^/1^L^) + log_2_(3^H^/1^L^) + log_2_(2^L^/1^H^) + log_2_(3^L^/1^H^)] where 1,2 and 3 denote Phactr1^XXX^, Phactr1^XXX^ΔC, and empty vector cells, and H and L denote samples from cells grown in R10K8 and R0K0 media respectively. Sequence logos were generated using WebLogo (https://weblogo.berkeley.edu/logo.cgi/).

The mass spectrometry proteomics data have been deposited to the ProteomeXchange Consortium (http://proteomecentral.proteomexchange.org) via the PRIDE partner repository with the dataset identifier PXD019977.

### TMT phosphoproteomics in neurons

The Phactr1-null (tm1d) allele was derived from the CSD79794 Phactr1 Tm1a allele (KOMP; https://www.komp.org/geneinfo.php?geneid=74176) by sequential action of Flp and Cre. Heterozygous animals were crossed. Hippocampal and cortical tissue was extracted from E16.5 embryos, and cultured in 12-well plate dishes (500,000 cells per well) as described (Baltussen et al., 2018) before genotyping. Two biological replicates were processed for wildtype or Phactr1-null neurons. On DIV10, neurons were treated with for 30 min with CD (10 μM) or LatB (1 μM) or vehicle (DMSO). Preparation of lysates, protease digestion and phosphopeptide enrichment was done as described with minor modifications (Eder et al., 2020). Phosphopeptides were eluted directly into the Orbitrap Fusion Lumos, operated with Xcalibur software, with measurement in MS2 and MS3 modes. The instrument was set up in data-dependent acquisition mode, with top 10 most abundant peptides selected for MS/MS by HCD fragmentation.

Raw mass spectrometric data were processed in MaxQuant (version 1.6.2.10); database search against the *Mus musculus* canonical sequences from UniProtKB was performed using the Andromeda search engine. Fixed modifications were set as Carbamidomethyl (C) and variable modifications set as Oxidation (M), Acetyl (Protein N-term) and Phospho (STY). The estimated false discovery rate was set to 1% at the peptide, protein, and site levels, with a maximum of two missed cleavages allowed. Reporter ion MS2 or Reporter ion MS3 was appropriately selected for each raw file.

Phosphorylation site tables were imported into Perseus (v1.6.1.2) for analysis. Contaminants and reverse peptides were cleaned up from the Phosphosites (STY) and the values normalised using Z-score function across columns.

Cortical and hippocampal as well as MS2/MS3 data across two biological replicates were pooled. (DMSO-CD) differences were calculated and compared between Phactr1 WT and KO neurons using the two-sample t-test. Phosphorylation sites exhibiting significantly different dephosphorylation in WT neurons compared with KO neurons were considered to be Phactr1-dependent. Sequence logos were generated using WebLogo (https://weblogo.berkeley.edu/logo.cgi/).

The mass spectrometry proteomics data have been deposited to the ProteomeXchange Consortium (http://proteomecentral.proteomexchange.org) via the PRIDE partner repository with the dataset identifier PXD019882.

## SUPPLEMENTARY FIGURE LEGENDS

**Figure S1.**
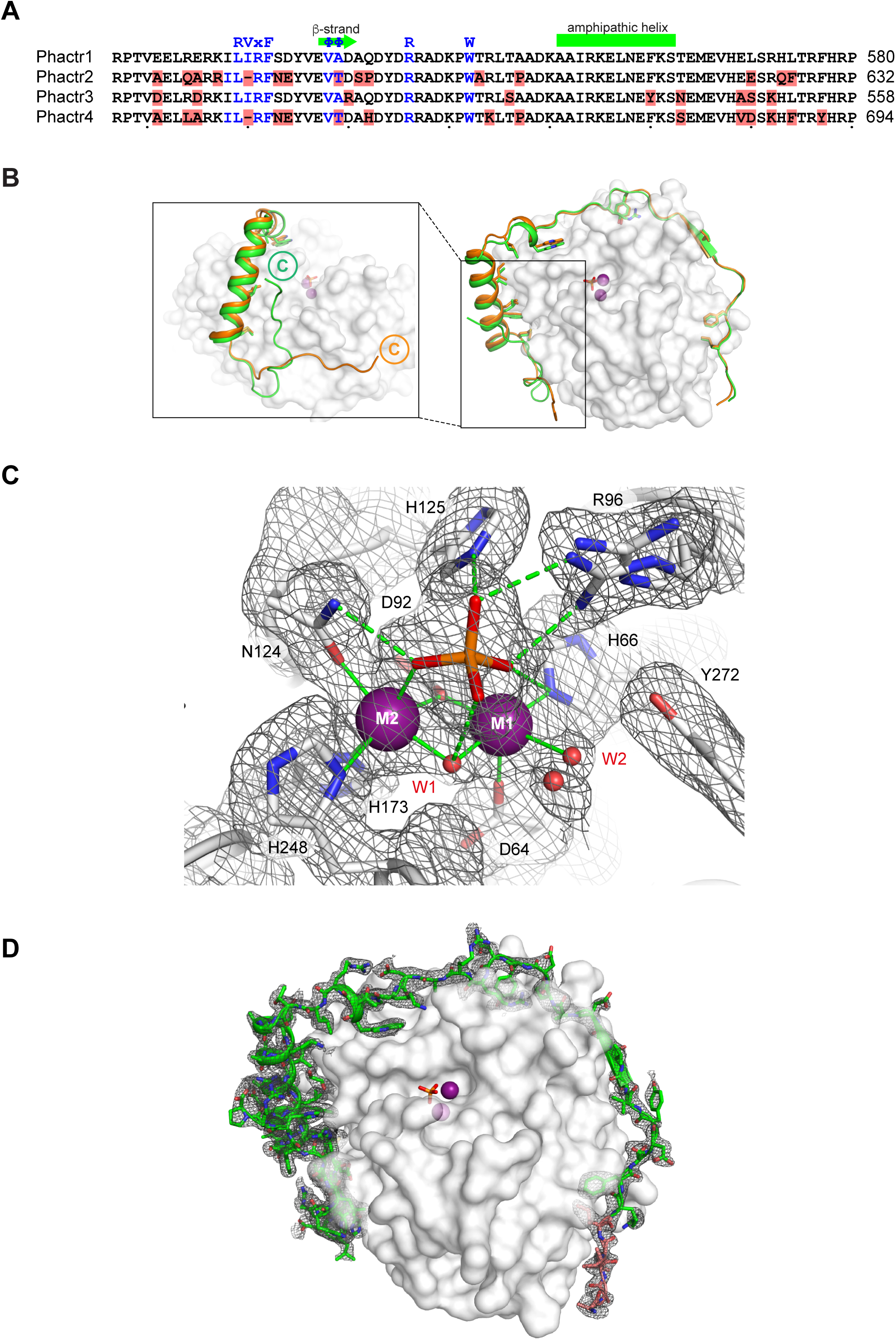
Crystallisation of Phactr1/PP1 complexes. (A) Sequence alignment of Phactr-family C-terminal sequences; differences are highlighted. (B) Structure of Phactr(507-580)/PP1α(7-300) solved at pH 8.5 (green) compared with that of Phactr1(516-580)/PP1α(7-300) solved at pH 5.25 (orange). The Phactr1 sequences C-terminal to the amphipathic α-helix (residue 467) adopt different conformations. In the pH 5.25 structure, residues 576-580 were not resolved in the density, and residues 567-575 were poorly resolved. (C) Omit map contoured at 3σ of the Phactr1/PP1 holoenzyme catalytic centre (metal ion/phosphate coordinating residues, white sticks; phosphate, orange sticks; water molecules, red spheres; coordination bonds, solid lines; hydrogen bonds, dashed lines. (D) Omit map contoured at 3σ of Phactr1 516-580 (green cartoon and sticks) across the PP1 surface (white).

**Figure S2.**
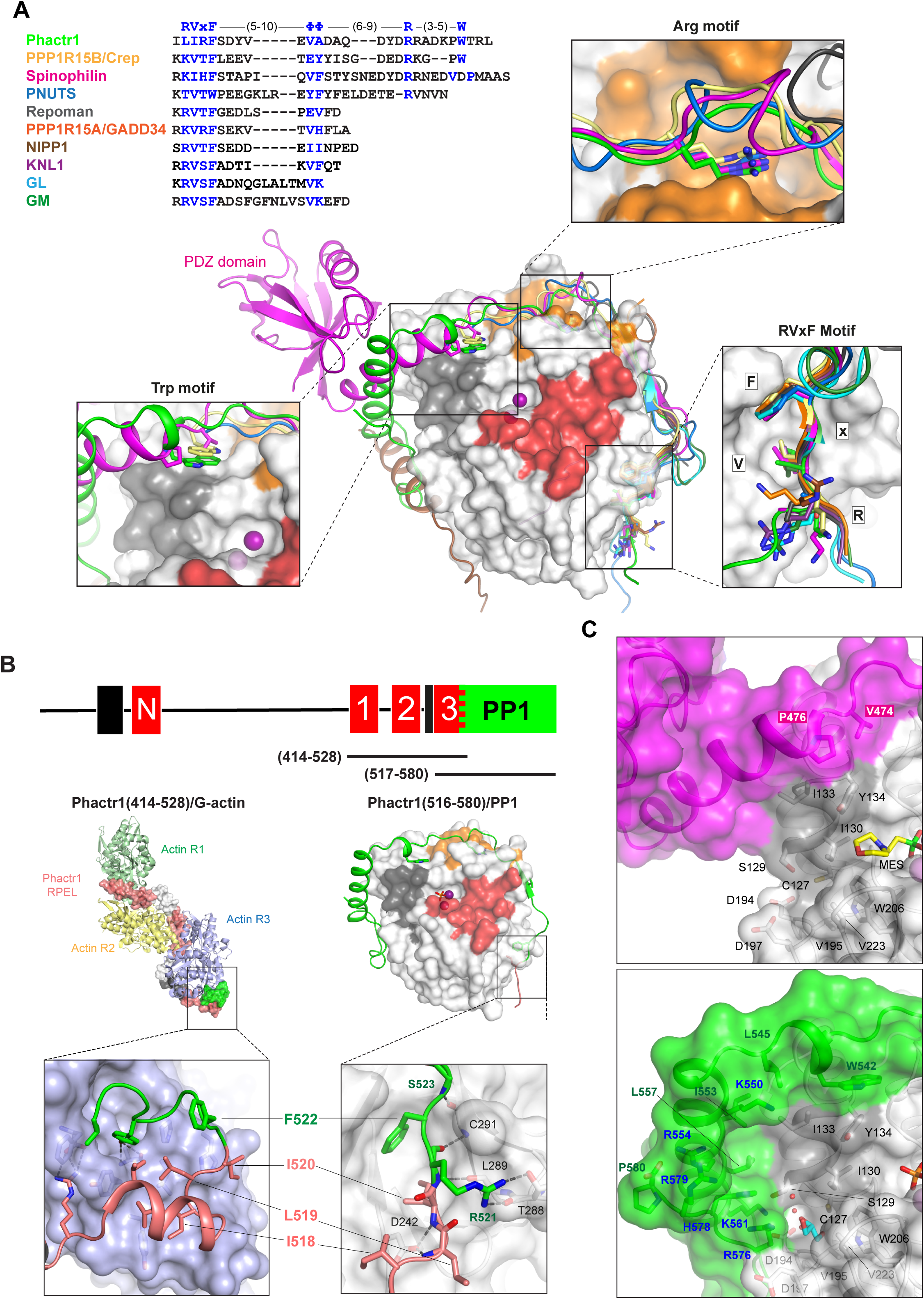
Phactr1 binds PP1 using an extended RVxF-ϕϕ-R motif. (A) Top, structure-based alignment of PIPs that bind PP1 through extended RVxF motifs including PPP1R15B (Chen et al., 2015), spinophilin (Ragusa et al., 2010), PNUTS (Choy et al., 2014), Repoman, (Kumar et al., 2016), PPP1R15A (Choy et al., 2015), NIPP1 (O’Connell et al., 2012), KNL1 (Bajaj et al., 2018), GL and GM (Yu et al., 2018). Below, comparison of the trajectories of Phactr1 (green), PPP1R15B (yellow), spinophilin (magenta; with PDZ domain), PNUTS (blue), Repoman (grey), PPP1R15A (orange), NIPP1 (brown), KNL1 (purple), GL (light blue) and GM dark green. Boxes show detail of interactions by the RVxF, Arg, and Trp motifs in each PIP. (B) Phactr1 RPEL3 (salmon) overlaps the RVxF motif LIRF(519-522) (green) and adopts different conformations when bound to G-actin (left) and PP1 (right). L519, I520 and F522 make critical but distinct hydrophobic contacts in both the G-actin and PP1 complexes. (C) Spinophilin remodels the PP1 hydrophobic groove in a different way to Phactr1. Top, the spinophilin/PP1 complex shown in surface representation (magenta, spinophilin; grey, PP1); bottom, the analogous region of the Phactr1/PP1 complex in the same orientation, with Figure 2B repeated for comparison).

**Figure S3.**
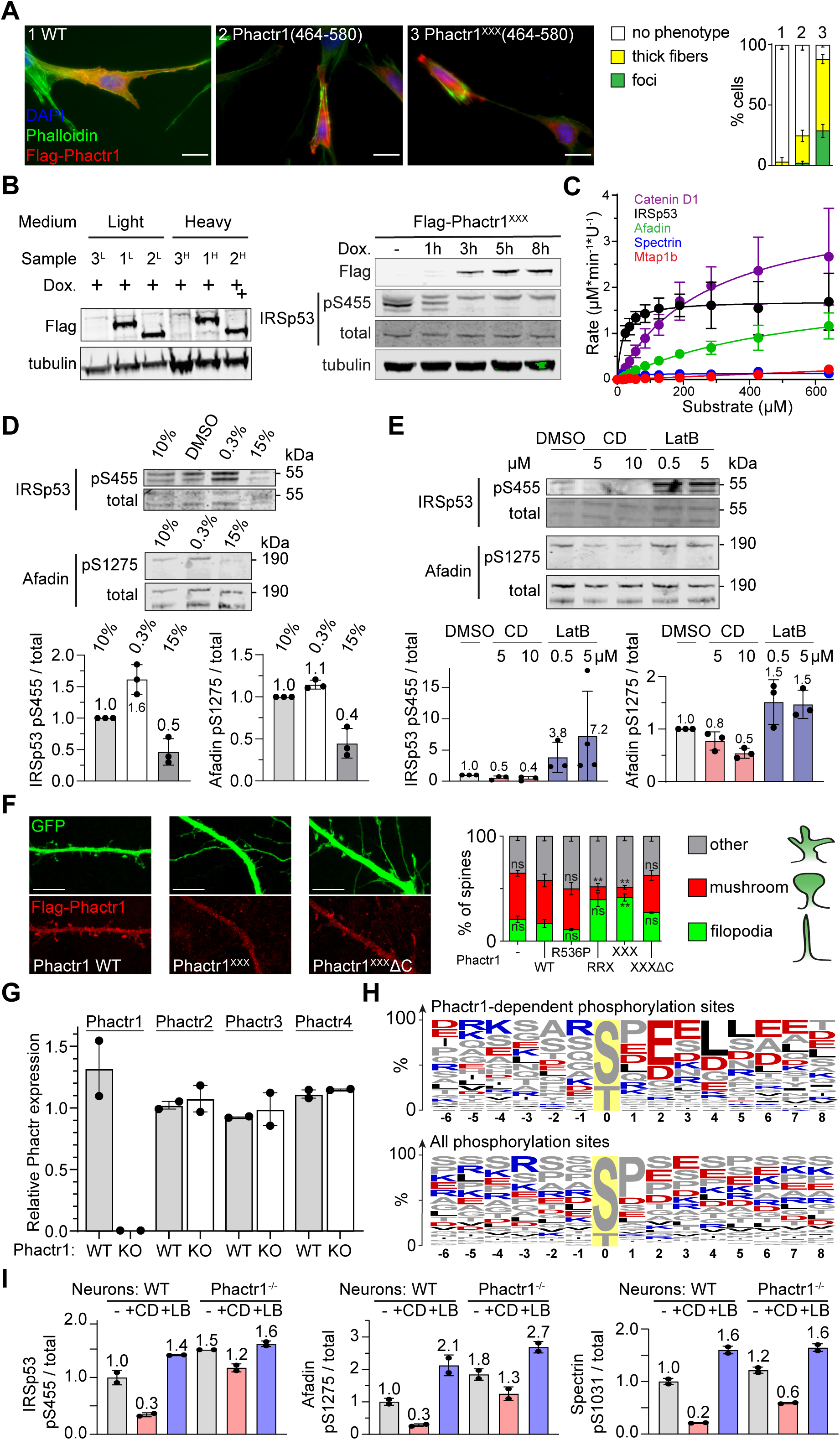
Substrates of the Phactr1/PP1 complex are enriched in cytoskeletal components and regulators. (A) Expression of the Phactr1^XXX^(464-580), which constitutively binds Phactr1, induces cytoskeletal rearrangements in NIH3T3 fibroblasts. Scalebar, 20 µm. (B) Left, immunoblot analysis of Flag-Phactr1^XXX^ and Flag-Phactr1^XXX^ΔC expression in NIH3T3 cell lines cultured under SILAC labelling conditions. Detection was with Flag antibody. Right, dephosphorylation of endogenous IRSp53 pS455 upon expression of Flag-Phactr1^XXX^, detected with anti-IRSp53 p455 antibody. For protein structures, see Figure 3A. (C). Phactr1/PP1 phosphatase activity assay data for selected substrates from Figure 3D (data are ± SD, n=3). (D, E) Immunoblot analysis of IRSp53 pS455 and afadin pS1275 in NIH3T3 cells upon overnight serum starvation and 30’ serum stimulation (D), or 30’ treatment with the actin-binding drugs cytochalasin D (CD) or latrunculin B (LB). Quantitation below (data are ± SD, n=3). (F) Rat hippocampal neurons were transfected with plasmids expressing Phactr1 derivatives and GFP, fixed after 1 day and their morphology scored blindly. Scalebar, 10 µm. Statistical significance was assessed by unpaired t-test with Welch’s correction (means ± SEM, n=3-5; **, p < 0.01). (G) RT-PCR analysis shows that Phactr1 inactivation in brain does not affect expression of other Phactr family members, relative to Rps16 expression. (H) Wildtype and Phactr1-null neurons were treated with cytochalasin D (CD) and phosphorylation changes assessed by TMT phosphoproteomics. Top, amino acid frequency table of phosphopeptides showing a statistically significant dependence on Phactr1 (FDR < 0.2), bottom, amino acid frequency table of all phosphopeptides. (I) Quantitation of immunoblot data shown in Figure 3F. Lysates from Phactr1-null or wildtype neurons were left untreated or treated with CD or LB for 30’ were analysed by immunoblotting with antibodies against IRSp53 pS455 (left); afadin pS1275 (centre); and spectrin αII pS1031 (right).

**Figure S4.**
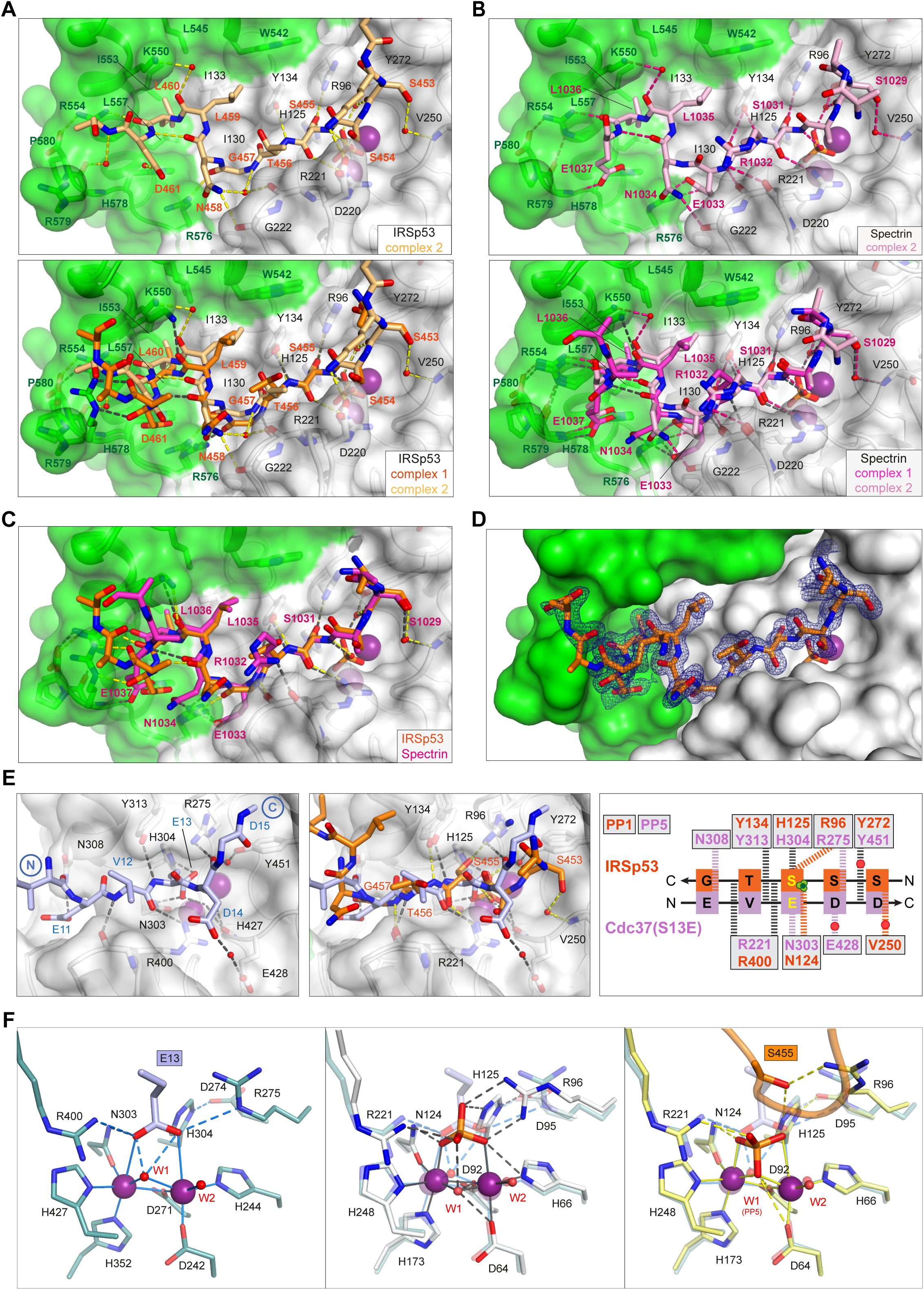
Substrate interactions in the Phactr1/PP1 complex. (A, B) Structure of the second copy of the Phactr1/PP1-substrate complexes in each asymmetric unit. Each structure is shown alone (top), and superimposed on the first copy (bottom; see Figure 4). (A) In IRSp53 complex 2, the electrostatic interactions between substrate residues +4, +5 and +6 and Phactr1 basic residues are water-bridged rather than direct. (B) In spectrin complex 2, the E1033/N1034^spectrin^ peptide bond is inverted, losing the mainchain carbonyl and N1034^spectrin^ sidechain interactions with Phactr1 R576, and the hydrogen bonding interaction between the L1035^spectrin^(+4) carbonyl and Phactr1 K550 is water-bridged rather than direct. (C) Superposition of the first copy of each complex (shown separately in Figure 4A, 4B). (D) Omit map contoured at 3σ of the first copy of the IRSp53 complex. (E) Opposite polarity of substrate binding to PP5 and Phactr1/PP1. Left: the PP5-Cdc37(S13E) complex (Oberoi et al., 2016). White surface, PP5; lilac sticks, Cdc37(S13E). Centre, superposition of the PP5-Cdc37(S13E) and Phactr1/PP1-IRSp53 structures. Right, comparison of molecular interactions at the catalytic site residues of PP5 (lilac) and PP1 (orange). Hydrogen bonds are shown as thick dashed lines: grey for both substrates; colour, for specific substrate. (F) Left, stick representation of molecular interactions at (left) the catalytic site of PP5-Cdc37(S13E). Centre, superposition of the PP5-Cdc37(S13E) (lilac) and Phactr1/PP1 complex (white) catalytic sites. Note the coincidence of the PP5-Cdc37(S13E) phosphomimetic glutamate with the oxygens of the phosphate present in the Phactr1/PP1 complex. Right, comparison of the PP5-Cdc37(S13E) and Phactr1/PP1-IRSp53 complexes (yellow). Note the inversion of the phosphate and the absence of W1.

**Figure S5.**
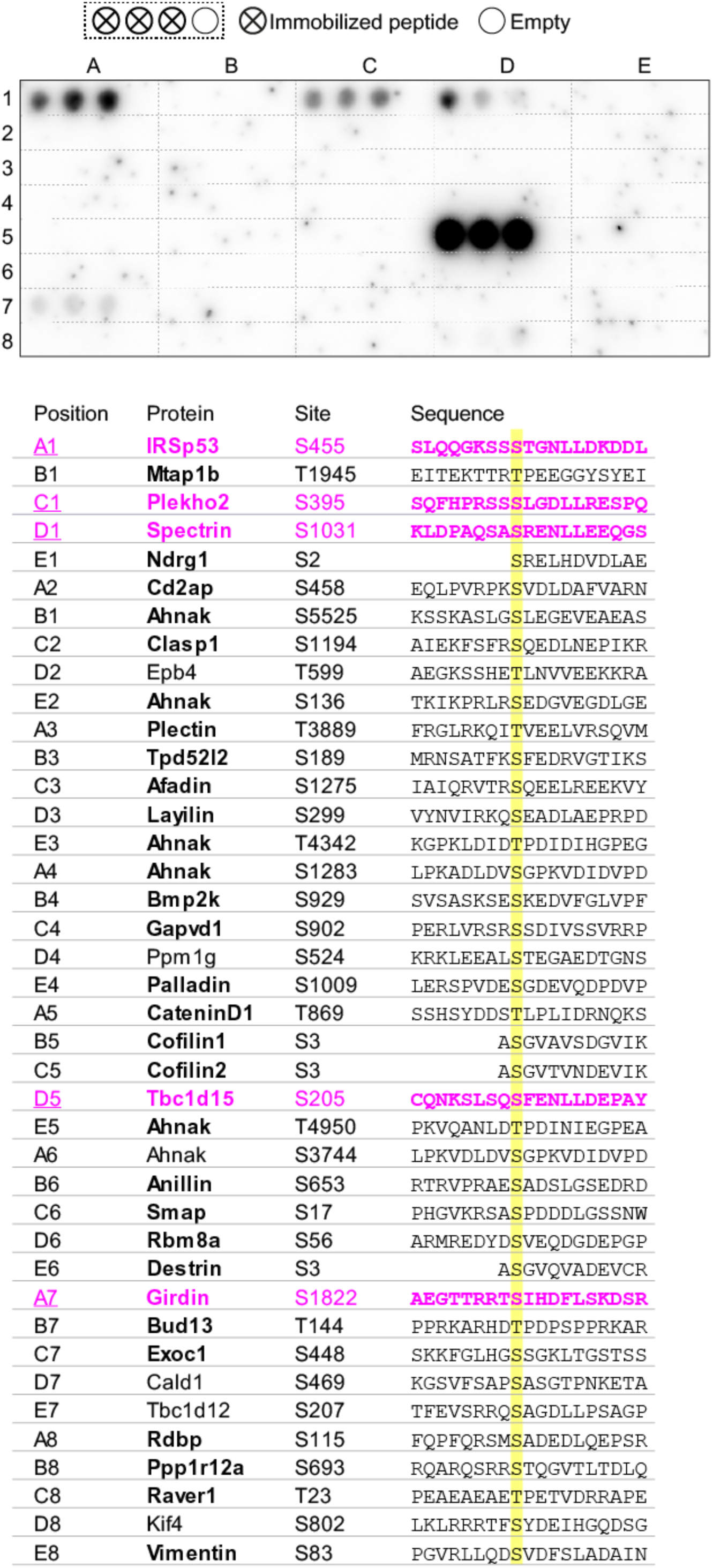
Binding of candidate Phactr1/PP1 substrates in the peptide array overlay assay. 19-mer unphosphorylated peptides from the indicated substrates were arrayed in triplicate as shown and tested for their ability to recruit GST-Phactr1(516-580)/PP1 from solution. Visualisation was with anti-GST antibody.

**Figure S6.**
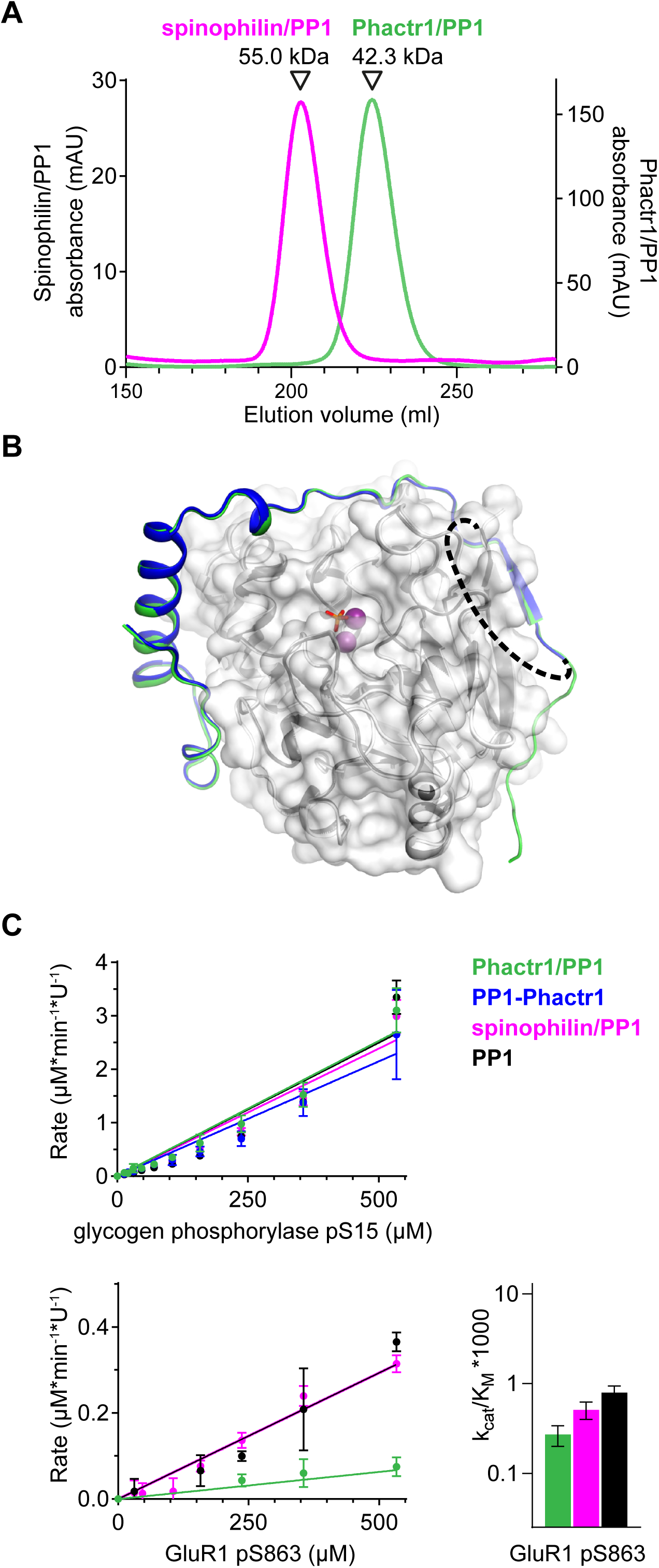
Flexible substrate interactions and substrate specificity of the Phactr1/PP1 complex. (A) Gel filtration profiles of the Phactr1/PP1 (green line) and spinophilin/PP1 complexes (magenta line) used for the substrate-specificity analysis. (B) Structure of the PP1-Phactr1 fusion protein, solved at 1.78 Å resolution. Phactr1 residues 526-580, dark blue cartoon; PP1, white surface; unresolved SGSGS linker, dashes; trajectory of Phactr1 residues 516-580 in the Phactr1/PP1 complex, green cartoon. PP1-Phactr1 exhibited 0.25 Å RMSD over 2395 atoms compared with the equivalent region of the Phactr1/PP1 holoenzyme complex. (C) Top, phosphatase activity data for 19mer peptides containing glycogen phosphorylase pS15 (top) and GluR1 pS863 (bottom). Relative catalytic efficiencies are shown in Figure 6E (glycogen phosphorylase pS15) or at right (GluR1 pS863) (data are ± half-range, n=2).

## SUPPLEMENTARY TABLES

**Table S1. Phosphoproteomics data**. (A, B) SILAC phosphoproteomics in NIH3T3 cells. (A) NIH3T3 cells expressing doxycycline-inducible Phactr1^XXX^, Phactr1^XXX^ΔC or vector alone were cultured in R0K0 or R10K8 SILAC media, protein expression induced with doxycycline, and cell lysates analysed by MS proteomics. Phosphorylation sites are annotated with dephosphorylation score, raw H/L values listed. (B) 1D enrichment analysis of Gene Ontology terms and kinase motifs based on the dataset from Table S1A. Terms with Benjamini-Hochberg FDR < 0.02 are shown. Gene Ontology Biological Process terms are in bold and those with positive mean value are reported in Figure 3B. (C-E) Wildtype (WT) or Phactr1-null (KO) cortical and hippocampal neurons DIV10 were treated with DMSO vehicle, CD or LB for 30’ and then analysed by TMT phosphoproteomics in MS2 and MS3 modes. Reporter ion intensities were normalized using Z-score function. (C) Raw and Z-score values for MS2 mode. (D) Raw and Z-score values for MS3 mode. (E) *t*-test was applied to (DMSO – CD) differences in WT neurons *vs* KO neurons. Phosphorylation sites are described and annotated with *t*-test difference and significance.

**Table S2. Peptides and Oligonucleotides**

**Table S3. Phosphatase Activity source data**

## Notes

https://www.rcsb.org

http://proteomecentral.proteomexchange.org

### Competing Interest Statement

The authors have declared no competing interest.

## REFERENCES

Adams, P.D., Afonine, P.V., Bunkoczi, G., Chen, V.B., Davis, I.W., Echols, N., Headd, J.J., Hung, L.W., Kapral, G.J., Grosse-Kunstleve, R.W., et al. (2010). PHENIX: a comprehensive Python-based system for macromolecular structure solution. Acta Crystallogr D Biol Crystallogr 66, 213–221.

Ali, S.R., Malone, T.J., Zhang, Y., Prechova, M., and Kaczmarek, L.K. (2020). Phactr1 regulates Slack (KCNT1) channels via protein phosphatase 1 (PP1). FASEB J 34, 1591–1601.

Allen, P.B., Greenfield, A.T., Svenningsson, P., Haspeslagh, D.C., and Greengard, P. (2004). Phactrs 1-4: A family of protein phosphatase 1 and actin regulatory proteins. Proc Natl Acad Sci U S A 101, 7187–7192.

Ba, W., Selten, M.M., van der Raadt, J., van Veen, H., Li, L.L., Benevento, M., Oudakker, A.R., Lasabuda, R.S.E., Letteboer, S.J., Roepman, R., et al. (2016). ARHGAP12 Functions as a Developmental Brake on Excitatory Synapse Function. Cell Rep 14, 1355–1368.

Bajaj, R., Bollen, M., Peti, W., and Page, R. (2018). KNL1 Binding to PP1 and Microtubules Is Mutually Exclusive. Structure 26, 1327–1336 e1324.

Baltussen, L.L., Negraes, P.D., Silvestre, M., Claxton, S., Moeskops, M., Christodoulou, E., Flynn, H.R., Snijders, A.P., Muotri, A.R., and Ultanir, S.K. (2018). Chemical genetic identification of CDKL5 substrates reveals its role in neuronal microtubule dynamics. The EMBO journal 37.

Bertran, M.T., Mouilleron, S., Zhou, Y., Bajaj, R., Uliana, F., Kumar, G.S., van Drogen, A., Lee, R., Banerjee, J.J., Hauri, S., et al. (2019). ASPP proteins discriminate between PP1 catalytic subunits through their SH3 domain and the PP1 C-tail. Nat Commun 10, 771.

Bobsin, K., and Kreienkamp, H.J. (2016). Severe learning deficits of IRSp53 mutant mice are caused by altered NMDA receptor-dependent signal transduction. J Neurochem 136, 752–763.

Bollen, M., Peti, W., Ragusa, M.J., and Beullens, M. (2010). The extended PP1 toolkit: designed to create specificity. Trends Biochem Sci 35, 450–458.

Bosch, M., Castro, J., Saneyoshi, T., Matsuno, H., Sur, M., and Hayashi, Y. (2014). Structural and molecular remodeling of dendritic spine substructures during long-term potentiation. Neuron 82, 444–459.

Boudrez, A., Beullens, M., Groenen, P., Van Eynde, A., Vulsteke, V., Jagiello, I., Murray, M., Krainer, A.R., Stalmans, W., and Bollen, M. (2000). NIPP1-mediated interaction of protein phosphatase-1 with CDC5L, a regulator of pre-mRNA splicing and mitotic entry. J Biol Chem 275, 25411–25417.

Brautigan, D.L., and Shenolikar, S. (2018). Protein Serine/Threonine Phosphatases: Keys to Unlocking Regulators and Substrates. Annu Rev Biochem 87, 921–964.

Chen, R., Rato, C., Yan, Y., Crespillo-Casado, A., Clarke, H.J., Harding, H.P., Marciniak, S.J., Read, R.J., and Ron, D. (2015). G-actin provides substrate-specificity to eukaryotic initiation factor 2alpha holophosphatases. Elife 4.

Choy, M.S., Hieke, M., Kumar, G.S., Lewis, G.R., Gonzalez-DeWhitt, K.R., Kessler, R.P., Stein, B.J., Hessenberger, M., Nairn, A.C., Peti, W., et al. (2014). Understanding the antagonism of retinoblastoma protein dephosphorylation by PNUTS provides insights into the PP1 regulatory code. Proc Natl Acad Sci U S A 111, 4097–4102.

Choy, M.S., Swingle, M., D’Arcy, B., Abney, K., Rusin, S.F., Kettenbach, A.N., Page, R., Honkanen, R.E., and Peti, W. (2017). PP1:Tautomycetin Complex Reveals a Path toward the Development of PP1-Specific Inhibitors. J Am Chem Soc 139, 17703–17706.

Choy, M.S., Yusoff, P., Lee, I.C., Newton, J.C., Goh, C.W., Page, R., Shenolikar, S., and Peti, W. (2015). Structural and Functional Analysis of the GADD34:PP1 eIF2alpha Phosphatase. Cell Rep 11, 1885–1891.

Cohen, D., Fernandez, D., Lazaro-Dieguez, F., and Musch, A. (2011). The serine/threonine kinase Par1b regulates epithelial lumen polarity via IRSp53-mediated cell-ECM signaling. J Cell Biol 192, 525–540.

Cohen, P.T. (2002). Protein phosphatase 1--targeted in many directions. J Cell Sci 115, 241–256.

Cox, J., and Mann, M. (2012). 1D and 2D annotation enrichment: a statistical method integrating quantitative proteomics with complementary high-throughput data. BMC Bioinformatics 13 Suppl 16, S12.

Crespillo-Casado, A., Claes, Z., Choy, M.S., Peti, W., Bollen, M., and Ron, D. (2018). A Sephin1-insensitive tripartite holophosphatase dephosphorylates translation initiation factor 2alpha. J Biol Chem 293, 7766–7776.

D’Arcy, A., Bergfors, T., Cowan-Jacob, S.W., and Marsh, M. (2014). Microseed matrix screening for optimization in protein crystallization: what have we learned? Acta Crystallogr F Struct Biol Commun 70, 1117–1126.

Diring, J., Mouilleron, S., McDonald, N.Q., and Treisman, R. (2019). RPEL-family rhoGAPs link Rac/Cdc42 GTP loading to G-actin availability. Nat Cell Biol 21, 845–855.

Eder, N., Roncaroli, F., Dolmart, M.C., Horswell, S., Andreiuolo, F., Flynn, H.R., Lopes, A.T., Claxton, S., Kilday, J.P., Collinson, L., et al. (2020). YAP1/TAZ drives ependymoma-like tumour formation in mice. Nat Commun 11, 2380.

Egloff, M.P., Cohen, P.T., Reinemer, P., and Barford, D. (1995). Crystal structure of the catalytic subunit of human protein phosphatase 1 and its complex with tungstate. J Mol Biol 254, 942–959.

Egloff, M.P., Johnson, D.F., Moorhead, G., Cohen, P.T., Cohen, P., and Barford, D. (1997). Structural basis for the recognition of regulatory subunits by the catalytic subunit of protein phosphatase 1. The EMBO journal 16, 1876–1887.

Emsley, P., Lohkamp, B., Scott, W.G., and Cowtan, K. (2010). Features and development of Coot. Acta Crystallogr D Biol Crystallogr 66, 486–501.

Fleming, M.R., Brown, M.R., Kronengold, J., Zhang, Y., Jenkins, D.P., Barcia, G., Nabbout, R., Bausch, A.E., Ruth, P., Lukowski, R., et al. (2016). Stimulation of Slack K(+) Channels Alters Mass at the Plasma Membrane by Triggering Dissociation of a Phosphatase-Regulatory Complex. Cell Rep 16, 2281–2288.

Goldberg, J., Huang, H.B., Kwon, Y.G., Greengard, P., Nairn, A.C., and Kuriyan, J. (1995). Three-dimensional structure of the catalytic subunit of protein serine/threonine phosphatase-1. Nature 376, 745–753.

Griffith, J.P., Kim, J.L., Kim, E.E., Sintchak, M.D., Thomson, J.A., Fitzgibbon, M.J., Fleming, M.A., Caron, P.R., Hsiao, K., and Navia, M.A. (1995). X-ray structure of calcineurin inhibited by the immunophilin-immunosuppressant FKBP12-FK506 complex. Cell 82, 507–522.

Hamada, N., Ogaya, S., Nakashima, M., Nishijo, T., Sugawara, Y., Iwamoto, I., Ito, H., Maki, Y., Shirai, K., Baba, S., et al. (2018). De novo PHACTR1 mutations in West syndrome and their pathophysiological effects. Brain 141, 3098–3114.

Hendrickx, A., Beullens, M., Ceulemans, H., Den Abt, T., Van Eynde, A., Nicolaescu, E., Lesage, B., and Bollen, M. (2009). Docking motif-guided mapping of the interactome of protein phosphatase-1. Chem Biol 16, 365–371.

Hirschi, A., Cecchini, M., Steinhardt, R.C., Schamber, M.R., Dick, F.A., and Rubin, S.M. (2010). An overlapping kinase and phosphatase docking site regulates activity of the retinoblastoma protein. Nat Struct Mol Biol 17, 1051–1057.

Hu, X.D., Huang, Q., Yang, X., and Xia, H. (2007). Differential regulation of AMPA receptor trafficking by neurabin-targeted synaptic protein phosphatase-1 in synaptic transmission and long-term depression in hippocampus. J Neurosci 27, 4674–4686.

Huet, G., Rajakyla, E.K., Viita, T., Skarp, K.P., Crivaro, M., Dopie, J., and Vartiainen, M.K. (2013). Actin-regulated feedback loop based on Phactr4, PP1 and cofilin maintains the actin monomer pool. J Cell Sci 126, 497–507.

Hurley, T.D., Yang, J., Zhang, L., Goodwin, K.D., Zou, Q., Cortese, M., Dunker, A.K., and DePaoli-Roach, A.A. (2007). Structural basis for regulation of protein phosphatase 1 by inhibitor-2. J Biol Chem 282, 28874–28883.

Ichikawa, K., Hirano, K., Ito, M., Tanaka, J., Nakano, T., and Hartshorne, D.J. (1996). Interactions and properties of smooth muscle myosin phosphatase. Biochemistry 35, 6313–6320.

Itoh, A., Uchiyama, A., Taniguchi, S., and Sagara, J. (2014). Phactr3/scapinin, a member of protein phosphatase 1 and actin regulator (phactr) family, interacts with the plasma membrane via basic and hydrophobic residues in the N-terminus. PLoS One 9, e113289.

Johnson, D.F., Moorhead, G., Caudwell, F.B., Cohen, P., Chen, Y.H., Chen, M.X., and Cohen, P.T. (1996). Identification of protein-phosphatase-1-binding domains on the glycogen and myofibrillar targetting subunits. Eur J Biochem 239, 317–325.

Kang, J., Park, H., and Kim, E. (2016). IRSp53/BAIAP2 in dendritic spine development, NMDA receptor regulation, and psychiatric disorders. Neuropharmacology 100, 27–39.

Kast, D.J., and Dominguez, R. (2019). Mechanism of IRSp53 inhibition by 14-3-3. Nat Commun 10, 483.

Kim, T.H., Goodman, J., Anderson, K.V., and Niswander, L. (2007). Phactr4 regulates neural tube and optic fissure closure by controlling PP1-, Rb-, and E2F1-regulated cell-cycle progression. Dev Cell 13, 87–102.

Kumar, G.S., Gokhan, E., De Munter, S., Bollen, M., Vagnarelli, P., Peti, W., and Page, R. (2016). The Ki-67 and RepoMan mitotic phosphatases assemble via an identical, yet novel mechanism. Elife 5.

Larson, J.R., Bharucha, J.P., Ceaser, S., Salamon, J., Richardson, C.J., Rivera, S.M., and Tatchell, K. (2008). Protein phosphatase type 1 directs chitin synthesis at the bud neck in Saccharomyces cerevisiae. Mol Biol Cell 19, 3040–3051.

Lei, W., Myers, K.R., Rui, Y., Hladyshau, S., Tsygankov, D., and Zheng, J.Q. (2017). Phosphoinositide-dependent enrichment of actin monomers in dendritic spines regulates synapse development and plasticity. J Cell Biol 216, 2551–2564.

Li, J., Wilkinson, B., Clementel, V.A., Hou, J., O’Dell, T.J., and Coba, M.P. (2016). Long-term potentiation modulates synaptic phosphorylation networks and reshapes the structure of the postsynaptic interactome. Science signaling 9, rs8.

Lin, C., Ear, J., Midde, K., Lopez-Sanchez, I., Aznar, N., Garcia-Marcos, M., Kufareva, I., Abagyan, R., and Ghosh, P. (2014). Structural basis for activation of trimeric Gi proteins by multiple growth factor receptors via GIV/Girdin. Mol Biol Cell 25, 3654–3671.

McCoy, A.J., Grosse-Kunstleve, R.W., Adams, P.D., Winn, M.D., Storoni, L.C., and Read, R.J. (2007). Phaser crystallographic software. J Appl Crystallogr 40, 658–674.

Miller, C.J., and Turk, B.E. (2018). Homing in: Mechanisms of Substrate Targeting by Protein Kinases. Trends Biochem Sci 43, 380–394.

Miralles, F., Posern, G., Zaromytidou, A.-I., and Treisman, R. (2003). Actin dynamics control SRF activity by regulation of its coactivator MAL. Cell 113, 329–342.

Mouilleron, S., Wiezlak, M., O’Reilly, N., Treisman, R., and McDonald, N.Q. (2012). Structures of the Phactr1 RPEL domain and RPEL motif complexes with G-actin reveal the molecular basis for actin binding cooperativity. Structure 20, 1960–1970.

Mueller, E.G., Crowder, M.W., Averill, B.A., and Knowles, J.R. (1993). Purple Acid Phosphatase: A Diiron Enzyme that Catalyzes a Direct Phospho Group Transfer to Water. J Am Chem Soc 115, 2974–2975.

O’Connell, N., Nichols, S.R., Heroes, E., Beullens, M., Bollen, M., Peti, W., and Page, R. (2012). The molecular basis for substrate specificity of the nuclear NIPP1:PP1 holoenzyme. Structure 20, 1746–1756.

Oberoi, J., Dunn, D.M., Woodford, M.R., Mariotti, L., Schulman, J., Bourboulia, D., Mollapour, M., and Vaughan, C.K. (2016). Structural and functional basis of protein phosphatase 5 substrate specificity. Proc Natl Acad Sci U S A 113, 9009–9014.

Pattison, M.J., Mitchell, O., Flynn, H.R., Chen, C.S., Yang, H.T., Ben-Addi, H., Boeing, S., Snijders, A.P., and Ley, S.C. (2016). TLR and TNF-R1 activation of the MKK3/MKK6-p38alpha axis in macrophages is mediated by TPL-2 kinase. Biochem J 473, 2845–2861.

Ragusa, M.J., Dancheck, B., Critton, D.A., Nairn, A.C., Page, R., and Peti, W. (2010). Spinophilin directs protein phosphatase 1 specificity by blocking substrate binding sites. Nat Struct Mol Biol 17, 459–464.

Robens, J.M., Yeow-Fong, L., Ng, E., Hall, C., and Manser, E. (2010). Regulation of IRSp53-dependent filopodial dynamics by antagonism between 14-3-3 binding and SH3-mediated localization. Mol Cell Biol 30, 829–844.

Sagara, J., Arata, T., and Taniguchi, S. (2009). Scapinin, the protein phosphatase 1 binding protein, enhances cell spreading and motility by interacting with the actin cytoskeleton. PLoS One 4, e4247.

Sagara, J., Higuchi, T., Hattori, Y., Moriya, M., Sarvotham, H., Shima, H., Shirato, H., Kikuchi, K., and Taniguchi, S. (2003). Scapinin, a putative protein phosphatase-1 regulatory subunit associated with the nuclear nonchromatin structure. J Biol Chem 278, 45611–45619.

Schrodinger, L. (2020). The PyMOL molecular graphics system, version 2.0.

Scita, G., Confalonieri, S., Lappalainen, P., and Suetsugu, S. (2008). IRSp53: crossing the road of membrane and actin dynamics in the formation of membrane protrusions. Trends Cell Biol 18, 52–60.

Swingle, M.R., Honkanen, R.E., and Ciszak, E.M. (2004). Structural basis for the catalytic activity of human serine/threonine protein phosphatase-5. J Biol Chem 279, 33992–33999.

Terrak, M., Kerff, F., Langsetmo, K., Tao, T., and Dominguez, R. (2004). Structural basis of protein phosphatase 1 regulation. Nature 429, 780–784.

Touati, S.A., Kataria, M., Jones, A.W., Snijders, A.P., and Uhlmann, F. (2018). Phosphoproteome dynamics during mitotic exit in budding yeast. The EMBO journal 37.

Vaguine, A.A., Richelle, J., and Wodak, S.J. (1999). SFCHECK: a unified set of procedures for evaluating the quality of macromolecular structure-factor data and their agreement with the atomic model. Acta Crystallogr D Biol Crystallogr 55, 191–205.

Vartiainen, M.K., Guettler, S., Larijani, B., and Treisman, R. (2007). Nuclear actin regulates dynamic subcellular localization and activity of the SRF cofactor MAL. Science 316, 1749–1752.

Wiezlak, M., Diring, J., Abella, J., Mouilleron, S., Way, M., McDonald, N.Q., and Treisman, R. (2012). G-actin regulates the shuttling and PP1 binding of the RPEL protein Phactr1 to control actomyosin assembly. J Cell Sci 125, 5860–5872.

Winter, G., Lobley, C.M., and Prince, S.M. (2013). Decision making in xia2. Acta Crystallogr D Biol Crystallogr 69, 1260–1273.

Yu, J., Deng, T., and Xiang, S. (2018). Structural basis for protein phosphatase 1 recruitment by glycogen-targeting subunits. FEBS J 285, 4646–4659.

Zhang, M., Yogesha, S.D., Mayfield, J.E., Gill, G.N., and Zhang, Y. (2013). Viewing serine/threonine protein phosphatases through the eyes of drug designers. FEBS J 280, 4739–4760.

Zhang, Y., Kim, T.H., and Niswander, L. (2012). Phactr4 regulates directional migration of enteric neural crest through PP1, integrin signaling, and cofilin activity. Genes & development 26, 69–81.

